# Local brain-age: A U-Net model

**DOI:** 10.1101/2021.01.26.428243

**Authors:** Sebastian G. Popescu, Ben Glocker, David J. Sharp, James H. Cole

## Abstract

We propose a new framework for estimating neuroimaging-derived “brain-age” at a local level within the brain, using deep learning. The local approach, contrary to existing global methods, provides spatial information on anatomical patterns of brain ageing. We trained a U-Net model using brain MRI scans from n=3463 healthy people (aged 18-90 years) to produce individualised 3D maps of brain-predicted age. When testing on n=692 healthy people, we found a median (across participant) mean absolute error (within participant) of 9.5 years. Performance was more accurate (MAE around 7 years) in the prefrontal cortex and periventricular areas. We also introduce a new voxelwise method to reduce the age-bias when predicting local brain-age “gaps”. To validate local brain-age predictions, we tested the model in people with mild cognitive impairment or dementia using data from OASIS3 (n=267). Different local brain-age patterns were evident between healthy controls and people with mild cognitive impairment or dementia, particularly in subcortical regions such as the accumbens, putamen, pallidum, hippocampus and amygdala. Comparing groups based on mean local brain-age over regions-of-interest resulted in large effects sizes, with Cohen’s *d* values >1.5, for example when comparing people with stable and progressive mild cognitive impairment. Our local brain-age framework has the potential to provide spatial information leading to a more mechanistic understanding of individual differences in patterns of brain ageing in health and disease.

## 1 Introduction

Brain ageing is associated with cognitive decline and an increased risk of neurodegenerative disease, though these effects vary greatly between individuals. Brain atrophy, often measured using structural MRI, is commonly seen in many neurological diseases [1, 2], but also in the normal ageing process. Even hippocampal atrophy, which is often thought to be characteristic of Alzheimer’s disease, can be seen in many other neurological and psychiatric conditions, and in normal ageing [3]. Evidently, both normal ageing and dementia can affect the same brain regions [4]. This fact complicates research into the earliest stages of age-related neurodegenerative diseases. Determining whether changes are ‘normal’ and or pathological is challenging. The brain-age paradigm can offer information on whether an individual’s brain is changing as expected for their age. The difference between chronological age and “brain-predicted age” obtained from neuroimaging data has been provided insights into the relationship between brain ageing and disease, and may be a useful biomarker for predicting clinical outcomes [5, 6, 7, 8]. For example, in Alzheimer’s Disease (AD), patients have previously been shown to have older-appearing brains, and that individuals with mild cognitive impairment (MCI) who had an older-appearing brain were more likely to progress to dementia [9, 10, 11, 12]. However, despite the growing literature employing the brain-age paradigm [13, 14], current approaches tend to generate brain-age predictions at a global level, with a single value per brain image. While some efforts have been made to derive patterns of ‘feature importance’ or similar from brain-age models [15, 16, 17, 18, 19], these patterns are at population-level, and do not apply to the individual.

### Localized Brain Predicted Age

Obtaining a finer-grained picture of brain-ageing patterns for a given brain disease is likely to provide several benefits. Firstly, neuroanatomical patterns should enable inferences to be made about mechanisms underlying the clinical manifestation of the disease. Secondly, better predictive discrimination between clinical groups should be possible, as different groups are likely to be associated with different spatial patterns of age-related brain changes, even in the case where ‘global’ brain-age differences are similar. Thirdly, the local individualised maps should enable fine-grain characterisation of brain changes over time, as the disease progresses or in response to treatment. Finally, spatial patterns of brain-age could be used to discover clinically-relevant subgroups in a data-driven manner, for example using clustering techniques.

### Related work

Limited prior work on local predictions of brain-age are available. Of note, is the early work of Cherubini et al. [20], who used linear regression models with voxel-level features derived from voxel-based morphometry and diffusion-tensor imaging to demonstrate reasonable prediction results in a small sample of healthy people (n=140). This approach of using a separate linear regression model for each voxel is limited as it does not incorporate contextual information from neighbouring voxels, and is insensitive to non-linear relationships. Other studies have provided local or regional information by training separate models per region e.g., [21], though again this precludes the incorporation of contextual and global information in the local predictions and is limited to the specific anatomical atlas used to define the brain regions.

Some studies have gone further and extracted ‘patch’ level information on brain-age, subsequently averaging predictions across brain regions to arrive at a global-level prediction [22, 23, 24, 25]. In Bintsi et al. [23], the authors use a ResNet [26] for each 64^3^ 3D block, reporting MAE values between 2.16 and 4.19 depending on block origin. While these approaches are promising, the relatively large size of the patch limits spatial resolution which results in less insightful inference in clinical settings. For example, semantic dementia is associated with a relatively localised spatial pattern of atrophy, often the left anterior and middle temporal lobe [27, 28], which could be overlooked by brain-age prediction models that lack spatial resolution. Alternatively, in Beheshti et al. [24], the authors introduce a model based on kernel methods introduced in [29], whereby they predict the grading at 7^3^ voxel patches. However, the authors use Support Vector Regression to aggregate the patch-level results to arrive at a global level prediction and do not provide patch-level results in the cortical regions. Similarly, [25] proposed a slice-level MRI encoding network, followed by an aggregation method to obtain global-level predictions. Likewise, the authors do not provide results at finer grained scales.

### Contributions

The goal of this work was to develop a model to accurately predict chronological age at the local level in healthy people, by incorporating voxelwise information using deep learning. U-Nets [30], which are typically used for tumor [31] or organ [32] segmentation, provide an excellent framework for voxelwise predictions, as their specific architecture enables the inclusion of contextual spatial information into individual predictions. Here, we introduce a deep learning algorithm that is trained to predict localised brain-age, producing high-resolution maps of brain-predicted age differences (brain-PAD maps) covering the entire brain (see Figure 1). We hypothesised that brain-PAD in healthy people would be centred on zero and would smoothly vary across regions of the brain. We further hypothesised that people with MCI and dementia patients would see higher brain-PAD values in regions previously reported to dementia-related atrophy. We provide an in-depth analysis of the structural differences seen in people with MCI and AD patients. We provide a means to reduce the so-called “age-bias” in brain-PAD maps and examine the reliability of local brain-age predictions, both within and between scanners.

**Figure 1:**
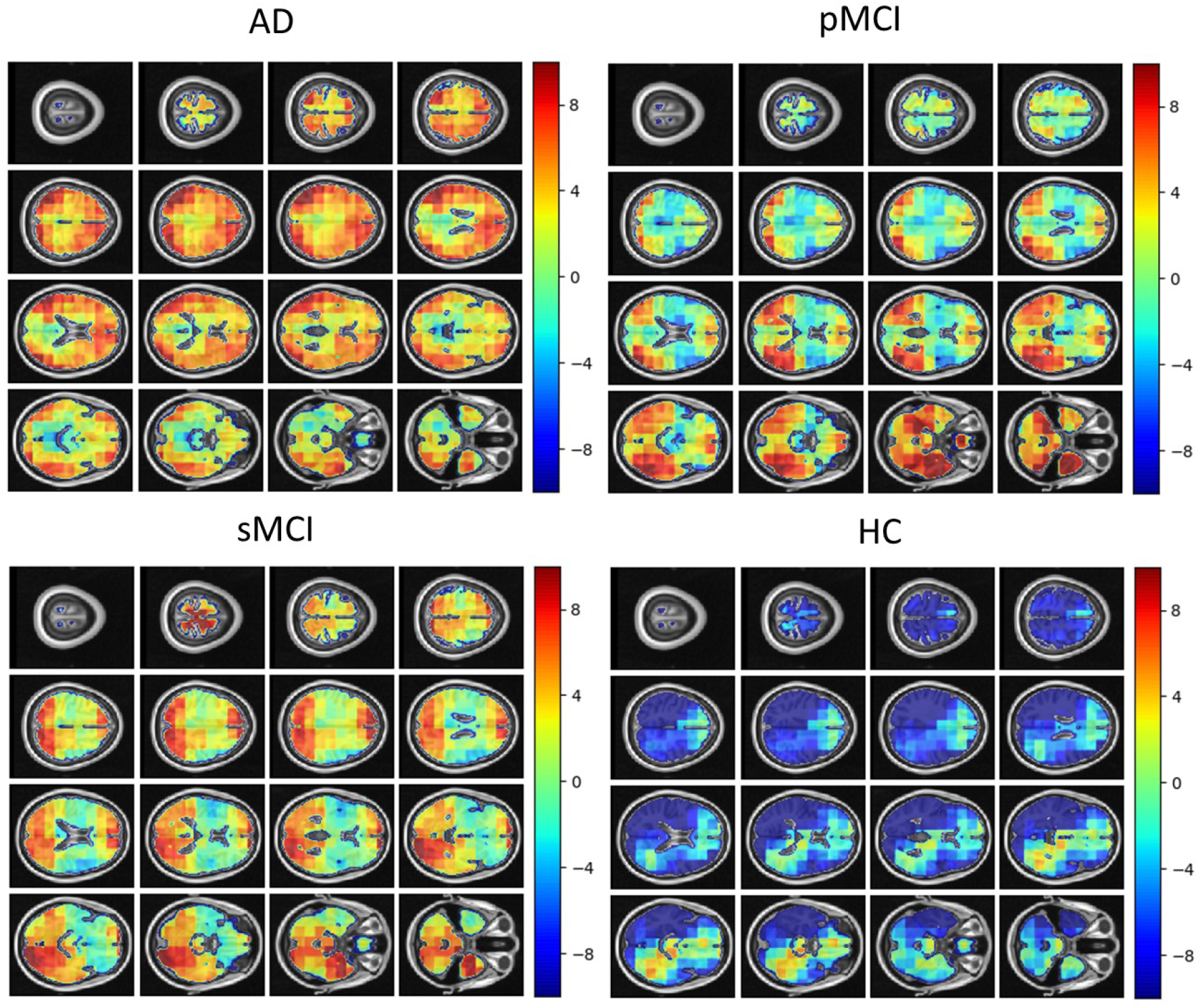
Local brain-PAD maps for randomly sampled participants from clinical groups in cross-sectional OASIS3 dataset. Positive values indicate an increased pattern of local volume differences compared to healthy ageing patterns at the respective age. HC = Healthy Controls, pMCI = progressive MCI, sMCI = stable MCI, AD = Alzheimer’s Disease.

## 2 Methods

### 2.1 Participants

To train, test and validate our local brain-age model, we collated multiple datasets comprising T1-weighted MRI brain scans. All included datasets were from studies that had been reviewed and approved by the local ethics committees and all participants provided informed consent. All participants were included, notwithstanding exclusions due to failure during quality control after pre-processing. All data were from publicly accessible databases. The supplementary material includes all links to access the respective databases, alongside chronological age histograms for each datasets (see Figure S1).

#### Brain-Age Healthy Controls (BAHC)

This dataset comprises 2001 healthy individuals with a male/female ratio of 1016/985, with a mean age of 36.95±18.12, aged 18-90 years. These data are an amalgam of 14 separate publicly-available datasets, as used in our previous brain-age research [33] (see supplementary material Table S1 for full details).

#### Dallas Lifespan Brain Study (DLBS)

This is a major effort designed to understand the antecedents of preservation and decline of cognitive function at different stages of the adult lifespan, with a particular interest in the early stages of a healthy brain’s march towards Alzheimer Disease. For our purpose we have selected solely the T1-weighted MRI scans, totaling n=315 healthy participants aged 18-89 years, with a mean age of 54.61±20.09 and male/female ratio of 117/198. All participants were scanned on a single 3T Philips Achieva scanner equipped with an 8-channel head coil. High-resolution anatomical images were collected with a sagittal T1-weighted 3D MP-RAGE sequence (TR = 8.1ms, TE = 3.7ms, flip angle = 12°, FOV = 204×256×160, slices = 160, voxel size = 1mm isotropic). More information can be found at https://dlbsdata.utdallas.edu/.

#### Cambridge Centre for Ageing and Neuroscience (Cam-CAN)

This dataset is part of larger project which is trying to use epidemiological, behavioural and neuroimaging data to understand how individuals can best retain cognitive abilities into old age. The dataset consists of n=652 T1-weighted MRI scans (3D MPRAGE, TR = 2250ms, TE = 2.99ms, TI = 900ms, flip angle = 9°, FOV = 256×240×192, voxel size = 1mm isotropic, GRAPPA=2) from participants aged 18-88 years, with a mean age of 54.29±18.59 and a male/female ratio of 322/330. More information can be found at https://www.cam-can.org/.

#### Southwest University Adult Lifespan Dataset (SALD)

This comprises a large cross-sectional sample (n = 494; age range = 19-80 years; mean age 45.18± 17.44; male/female ratio of 187/307) undergoing a multi-modal (structural MRI, resting state fMRI, and behavioral) neuroimaging. Only T1-weighted MRI (3D MPRAGE, TR = 1900ms, TE = 2.52ms, TI = 900ms, flip angle = 90°, matrix = 256×256, slices = 176, voxel size = 1mm isotropic) were used here. The goals of the SALD are to give researchers the opportunity to map the structural and functional changes the human brain undergoes throughout adulthood and to replicate previous findings. More information can be found at http://fcon_1000.projects.nitrc.org/indi/retro/sald.html.

#### Wayne State

The Wayne State longitudinal data set for the Brain Aging in Detroit Longitudinal Study, comprises 200 healthy individuals, with n=302 total anatomical scans across two waves of data collection and mean age of 53.94±15.58, with a male/female ratio of 37/77. All the participants were screened by the local research centres to be free from neurological or psychiatric disorders according to well established protocols. All of the neuroimaging data were acquired either at 1.5T or 3T using standard T1-weighted sequences (TR = 8000ms, TE = 3.93ms, TI = 420ms, flip angle = 20°), FOV = 256×192×100 averages = 3, voxel size = 0.75×0.075×1.5mm. More information can be found at http://fcon_1000.projects.nitrc.org/indi/retro/wayne_10.html.

#### Within-scanner reliability dataset

Here we used data from the Imperial College London project, STudy Of Reliability of MRI (STORM). The study comprises of 20 participants with a male-female ratio of 12/8, with a mean age at the first scan undertaken of 34.05 ± 8.71. The participants were scanned for the second time at an average distance of 28.35 ± 1.09 days. All participants were free from any neurological or psychiatric disorders. T1-weighted MRI data were acquired using a Siemens Verio 3T scanner (3D MPRAGE, FOV = 240×256×160, voxel size = 1mm isotropic).

#### Scanner calibration dataset

This study included 11 participants scanned in two different centres, mean age at first scan of 30.88 ± 6.16 and with a male/female ration of 7/4. The two scanning sites were at Imperial College London, where a Siemens Verio 3T scanner was used (3D MPRAGE, FOV = 240×256×160, voxel size = 1mm isotropic), whereas a Philips Ingenia 3T scanner was used at the Academic Medical Center Amsterdam (T1-TFE, FOV = 256×256×170, voxel size = 1.05×1.05×1.2mm). The mean interval between scans was 68.17±92.23 days.

#### Open Access Series of Imaging Studies (OASIS3)

This is a retrospective compilation of data for >1000 participants that were collected across several ongoing projects through the WUSTL Knight ADRC over the course of 30 years. Participants include n=609 cognitively normal adults and n=489 individuals at with MCI or dementia ranging in age from 42-95 years. Using Clinical Dementia Rating scale (CDR) scores, we classified participants as healthy control (HC), stable MCI, progressive MCI or AD, as detailed in Table 1. Follow-up CDR scores used to define MCI status were from at least 3 years after baseline assessments. We excluded scans which did not pass quality standards after pre-processing pipeline. MRI was collected on 3 different Siemens scanner models: Vision 1.5T, TIM Trio 3T and BioGraph mMR PET-MR 3T. Further information can be found at https://www.oasis-brains.org/.

**Table 1:** Demographic characteristics for the OASIS3 dataset. CDR = Clinical Dementia Rating scale, HC = Healthy Controls, pMCI = progressive MCI, sMCI = stable MCI, AD = Alzheimer’s Disease.

| Characteristics | HC<br>(n=128) | sMCI<br>(n=29) | pMCI<br>(n=29) | AD<br>(n=78) |
| --- | --- | --- | --- | --- |
| Males/Females, n | 70/58 | 15/14 | 18/11 | 33/45 |
| Age, mean (SD) years | 68.14 (9.40) | 76.44 (6.81) | 75.72 (7.68) | 75.02 (8.90) |
| Age, range years | 42.66-97.11 | 59.2-94.44 | 49.38-93.93 | 50.35-95.58 |
| Baseline CDR | 0.0 | 0.5 | 0.5 | $\geq 1.0$ |
| Follow-up CDR | 0.0 | 0.5 | $\geq 1.0$ | - |

#### Australian Imaging Biomarkers and Lifestyle Study of Aging (AIBL)

This study is a study to discover which biomarkers, cognitive characteristics, and health and lifestyle factors determine subsequent development of symptomatic Alzheimer’s Disease (AD) (https://aibl.csiro.au/). The dataset contained n=198 participants with Clinical Dementia Rating scale scores, detailed in Table 2.

**Table 2:** Demographic characteristics for the AIBL dataset. CDR = Clinical Dementia Rating scale, HC = Healthy Controls, pMCI = progressive MCI, sMCI = stable MCI, AD = Alzheimer’s Disease.

| Characteristics | HC<br>(n=83) | sMCI<br>(n=64) | pMCI<br>(n=20) | AD<br>(n=31) |
| --- | --- | --- | --- | --- |
| Males/Females, n | 38/45 | 36/26 | 11/9 | 17/14 |
| Age, mean (SD) years | 67.28 (9.70) | 75.83 (8.85) | 71.86 (9.21) | 74.23 (10.21) |
| Age, range years | 60.11-86.45 | 62.54-87.86 | 54.66-81.24 | 61.75-87.89 |
| Baseline CDR | 0.0 | 0.5 | 0.5 | $\geq 1.0$ |
| Follow-up CDR | 0.0 | 0.5 | $\geq 1.0$ | - |

### 2.2 Data pre-processing

All T1-weighted brain MRI scans were pre-processed using the Statistical Parametric Mapping (SPM12) software package (https://www.fil.ion.ucl.ac.uk/spm/software/spm12/). This entailed tissue segmentation into grey matter (GM) and white matter (WM), followed by a nonlinear registration procedure using the DARTEL algorithm [34] to the Montreal Neurological Institute 152 (MNI152) space, subsequently followed by resampling to 1.5*mm*^3^ with a 4mm smoothing kernel.

### 2.3 Statistical analysis

#### Inferential statistics

Welch’s t-test was used to compare groups based on voxel, regional and global brain-PAD values. Welch’s t-test is an alternative to the standard student’s t-test when the two populations to be compared have uneven variance and optionally also uneven sample size. The t statistics to test whether the populations means is given by:

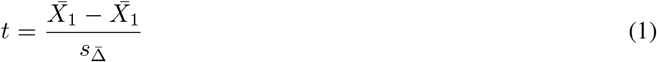

where 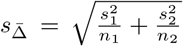 and 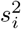 represents the unbiased estimator of the variance of a respective sample with *n_i_* participants. To use the test statistics for significance testing, the degrees of freedom of the associated Student’s t-distribution is given by the Welch-Satterthwaite equation:

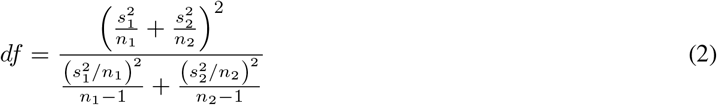

#### Effect size estimates

To quantify effect sizes when comparing different disease groups we used the standardised effect size Cohen’s *d*:

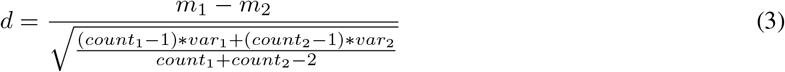

where *m_k_* is the mean, *var_k_* represents the variance, whereas *count_k_* defines the number of participants within group *k*. The purpose of this method is to quantify the size of the difference, allowing us to decide if the difference is meaningful.

#### Intraclass Correlation Coefficient

The intraclass correlation coefficient (ICC) is used to test the reproducibility of a certain quantitative measurement made by a specified number of observers which rate the same participant. The original formula is given as follows:

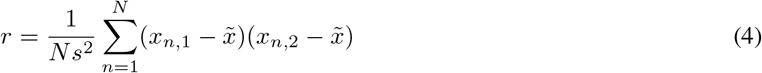

where 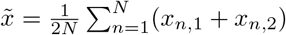 and 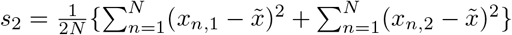

Here, we used ICC[2,1] as defined by Shrout and Fleiss [35]. The interval of values ranges from [−1,1] with values closer to 1 denoting that the observers (e.g., MRI scans or scanners) agree with each other.

### 2.4 Study design

In this subsection we summarize the design of our experiments and which datasets are used in each step.

- BAHC, CamCan, Dallas and SALD were used as training and validation sets (80/20 split) for training the local brain-age U-net model.
- All Wayne State participants and healthy participants from OASIS3 and AIBL were used at testing time. We calculated mean absolute error (MAE) values globally (by averaging across voxel-level brain-predicted age for each participant) and at voxel level, alongside the Pearson’s correlation coefficient between chronological age brain-predicted age at both global and voxel level.
- Within-scanner reliability and Scanner calibration datasets were used as test sets to compute voxel-level ICC values to assess the reliability of local brain-age when the same participant is scanned in two different scanners, respectively one the same scanner with short time interval.
- Using subcortical and cortical ROIs from the Harvard-Oxford structural brain atlas, we obtained brain tissues volumes (mm^3^) for each ROI, alongside ROI-level brain-PAD values which were computed by first averaging voxel-level “brain-predicted age” inside an ROI, then subtracting the participant’s chronological age. We calculated a Pearson’s correlation coefficient for each ROI.
- OASIS3 was used at testing time to assess the sensitivity of local brain-age to differences in brain structure between groups. For each participant we computed an mean across voxels global brain-PAD (adjusted for age bias), which we use then to perform Welch’s t-test between disease groups, correcting for multiple comparisons using the Bonferroni method. The same method was then applied to local-level brain-PAD values, which were pooled to create a “population” of brain-PAD values at voxel-level. To assess effect sizes of between-group differences we calculated Cohen’s *d* coefficient at global and voxel levels. Lastly, to assess regional differences between disease groups, we used the Harvard-Oxford cortical and subcortical structural atlas, which contains 48 cortical and 21 subcortical structural ROI. We then calculated differences in mean local brain-PAD per ROI between groups using Welch’s t-test (Bonferroni corrected), again computing Cohen’s *d* effect sizes. For all experiments in this part we have selected subjects above 60 years old from the healthy controls so as to have a similar chronological age distribution in relation to the groups with varying degrees of cognitive impairment.

A visual overview of the study design is portrayed in Figure 2.

**Figure 2:**
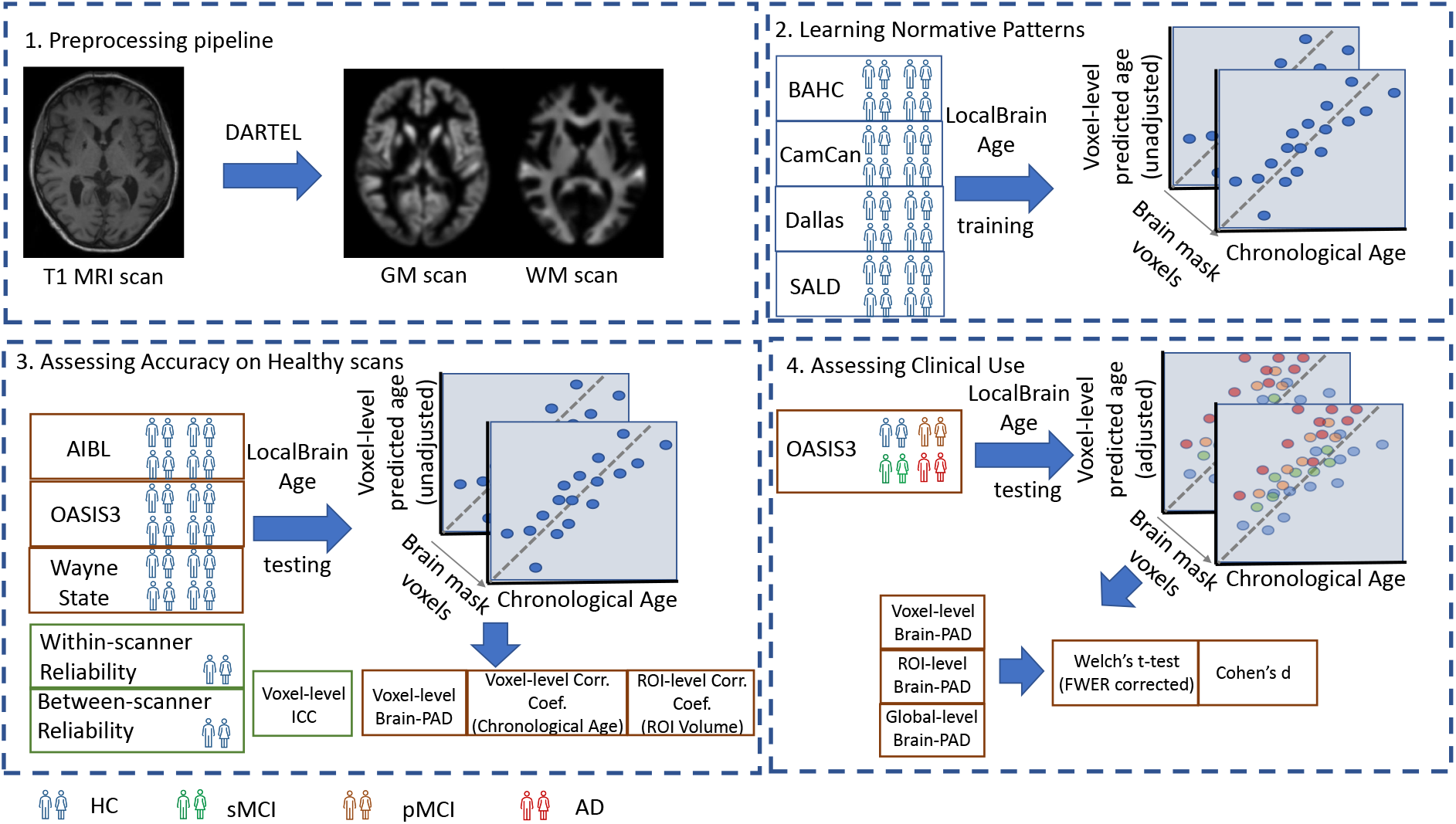
**1. Preprocessing pipeline**: T1-weighted MRI scans from all datasets were tissue segmented and non-linearly registered using SPM12 to generate modulated grey and white matter volume maps. **2. Learning Normative Patterns**: Randomly subsampled participants from BAHC, CamCan, Dallas and SALD (80%/20% split training/validation set) were used to train the local brain-age U-net to learn predict chronological age locally. **3. Assessing Accuracy on Healthy scans**: Using healthy scans from AIBL, OASIS3 and Wayne State we tested age prediction accuracy on unseen data from independent datasets. We calculated descriptive statistics for voxel-level brain-PAD, alongside Pearson’s correlation coefficients between chronological age and voxel-level “brain-ages” and between brain tissues volume and ROI-level “brain-ages” predictions using subcortical and cortical ROIs from the Harvard-Oxford atlas. To assess the reliability of local brain-age predictions with respect to between-scanner and within-scanner differences, we used the Within-scanner reliability dataset and Scanner calibration dataset, computing voxel-level Intraclass Correlation Coefficients (ICC). **4. Assessing Clinical Use**: Using the OASIS3 dataset (healthy controls, stable MCI, progressive MCI and AD participants), we compared groups using Welch’s t-test (family-wise error rate corrected), and calculated Cohen’s *d* effect sizes at the voxel, regional and global levels.

### 2.5 Local “Brain-age” Prediction

We used a fully convolutional neural network (CNN) inspired by the U-Net architecture introduced in Ronneberger et al. [30]. Our network architecture is illustrated in Figure 3. Input images were the output from SPM12 pre-processing, representing voxelwise maps of GM and WM volume. These images were split into overlapping 3-dimensional blocks of size 52^3^ voxels. The convolutional layers in our network used an isotropic 3×3×3 filter, convolved over the input image after which element-wise multiplication with the filter weights and subsequent summation was performed at each location. Subsequently, to allow for non-linear modelling, we passed the obtained values through an “activation function”; we used a LeakyReLu with alpha=0.2. *LeakyReLu*(*α*) are defined by the following equation *LeakyReLu*(*x*) = *max*(*x*, 0) + *min*(*x* * *α*, 0), thus allowing a small, non-zero gradient when the unit is not active.

**Figure 3:**
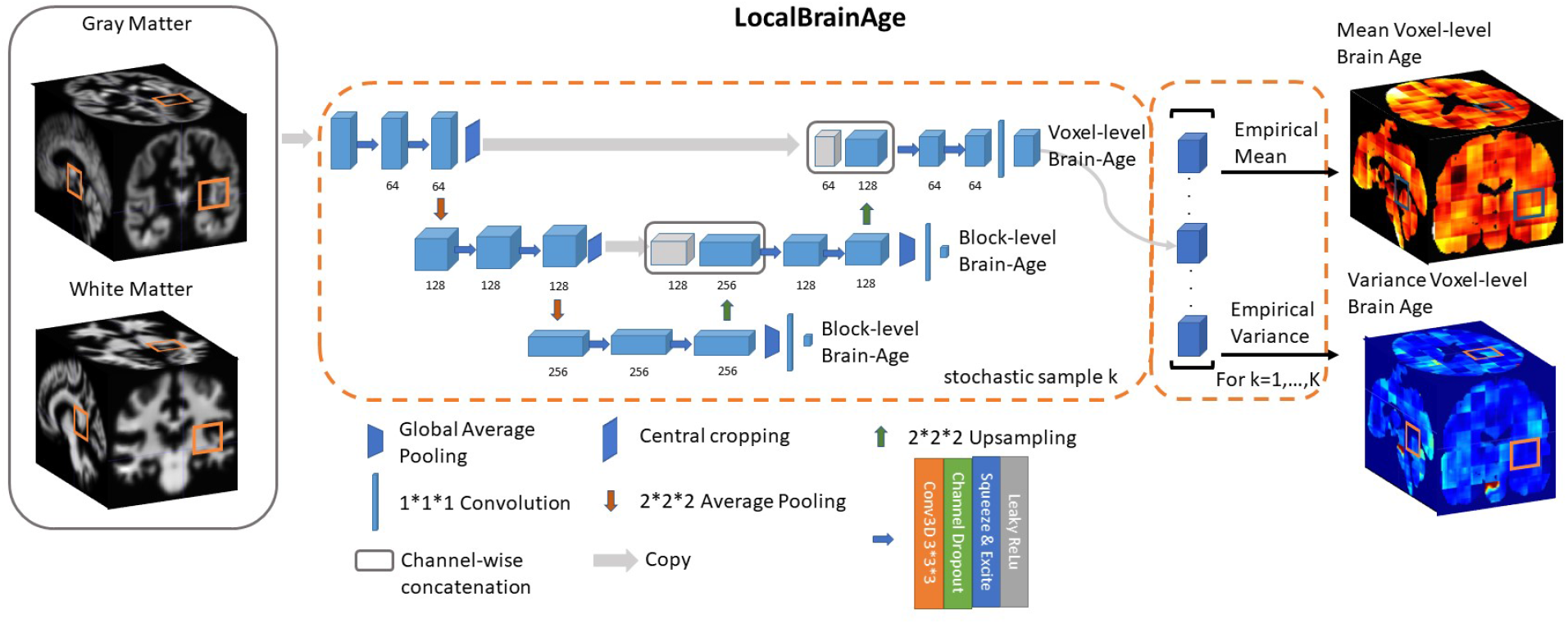
U-Net architecture for voxel-level brain-age prediction. Raw T1-weighted MRI scans were pre-processed using SPM12, obtaining modulated grey and white matter volume maps registered to the MNI152 template. Additional auxiliary block-level brain-age loss functions were added at each level of the U-Net to facilitate training.

The convolution operation is also controlled by its stride, which is how many pixels/voxels are skipped after every element-wise weight multiplication and summation. We set the stride equal to 1.

Downsampling increases the effective field of view or “receptive field” of layers higher in the hierarchy. For the downsampling part of the U-Net we used at each scale two consecutive 3D 3×3×3 filter kernels with an initial number of channels = 64, which get multiplied by 2 as we progress down the downsampling path. For downsampling we used 2×2×2 average pooling.

For the upsampling part of the U-Net we inverted the downsampling architecture, with the downsampling layers being replaced by 2×2×2 upsampling layers. At each convolution we used a squeeze-and-excite unit. Squeeze & Excite networks were introduced in Hu et al. [36] and can be viewed as computationally less intensive method of performing attention over the channels of a given feature block. Finally, at the end of the network we obtain predictions over 12^3^ voxels blocks.

Besides the voxel-level mean absolute error cost function on the output layer we introduced two additional cost functions at the two other scales of the architecture. We applied global average pooling followed by a dense layer to predict brain-age at block-level. The loss function can be expressed as follows:

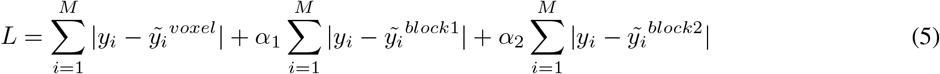

During training, we observed that the addition of these auxiliary loss functions helped stabilise the learning process. During training, *α*_1_ and *α*_2_ are progressively decreased so that the gradients will exclusively flow from the voxel-level predictions after 50,000 training iterations. We used Adam [37] for optimizing our loss function with a learning rate set to 0.0001. We trained our model for 500,000 iterations, with a minibatch size of 32 (gradient averaging over four splits). We split our healthy participant datasets into training (80%) and validation (20%) sets and the stopping criteria was set based on a visual inspection of the validation loss reaching a plateau. The model was implemented in Tensorflow [38].

### 2.6 Removing bias in predictions

Subtracting chronological age from estimated brain age provides a measure of the difference between an individual’s predicted age and chronological age, also known as the brain-age ‘gap’, brain-predicted age difference (brain-PAD) or brain-age ‘delta’. A so-called ‘regression dilution’ has been commonly observed in brain-age prediction algorithms, caused by noise in the neuroimaging features leading to a greater under- or over-estimate of age, the further away a sample is from the training set mean age. In other words, this effect results in the systematic under-estimation of brain-predicted age for older participants and over-estimation for younger participants, which increases as model performance decreases.

#### 2.6.1 Global-level

Broadly speaking, two approaches to account for this effect have been reported:

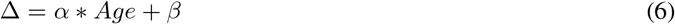

where Δ is the brain-age delta of a group of participants from an external dataset that is used specifically for adjusting the bias. *α* and *β* are the parameters of a linear regression with the covariate *Age* representing chronological age.

Then, to obtain the bias-adjusted age we have the following equation:

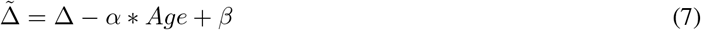

Another approach involves using the brain-predicted age in the linear regression, more specifically:

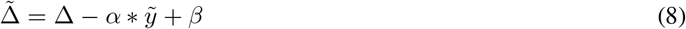

where 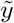 denotes the brain-predicted age. de Lange and Cole [39] showed that using either formulation results in the same statistical outcome in comparing different disease groups. The authors also argue against using bias adjusted predictions at testing time to assess overall accuracy of the model. However, this standard method used for global-level brain-age prediction did not succeed in de-biasing our predictions at voxel-level. Additional results using this approach are shown in the supplementary material (see Figure S2).

#### 2.6.2 Voxel-level

Here, we used a separate small batch (n=200) of participants randomly selected from the healthy participant datasets (BAHC, CamCan, Dallas, SALD), who were not included in the training or validation set. We obtained testing time predictions for these participants and calculated their voxel-level brain-age delta Δ*_i,v_*, where *i* indicates the i-th participant and *v* the v-th voxel. We then binned these participants based on their chronological age (5-year intervals, expect the first being between 18-25 years). Then, for each bin *b* we calculated the average voxel-level brain-age delta for that respective bin, which we denote as Δ*_b,v_*. This value will represents the average brain-age delta for that voxel given the chronological age interval. Subsequently, to de-bias the voxel-level brain-age delta for a new participant (e.g., from testing set), Δ*_j,v_* we used the following formula:

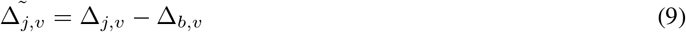

This method was used subsequent analysis where indicated.

## 3 Results

### 3.1 Local brain-age model performance in independent healthy test datasets

We tested the local brain-age model on healthy participants combined from the OASIS3 (n = 128), AIBL (n = 83) and Wayne State (n = 200) datasets. Individual local brain-age maps from example participants are shown in Figure 1. When looking at voxel-level MAE (unadjusted) values across the brain mask (Figure 4a), we mean values for AIBL of 10.84 ± 2.05 (median 10.43) years, for Wayne State 9.28 ± 1.05 (median 9.08) years, and OASIS3 9.70 ± 1.53 (median 9.33) years. The voxel-level MAE (unadjusted) values of the model varied in different brain regions. We observed lower values across the different sites in the prefrontal cortex and subcortical regions and higher MAE in the occipital lobe, cerebellum and brainstem (Figure 5a). The correlation coefficient between chronological age and voxel-level predicted brain-age (unadjusted) across participants showed similar patterns, with higher values obtained in the prefrontal cortex and subcortical regions (Figure 5b). We obtained a global-level MAE (unadjusted) value by averaging the voxel-level brain-predicted ages across voxels for a given participant. For AIBL, we obtain an average of 10.23 ± 7.08 years (median = 8.86; *r* = 0.47), for Wayne State 8.09 ± 6.08 years (median = 6.92; *r* = 0.78), respectively for OASIS3 we get an average of 8.08 ± 6.40 years (median = 6.26; *r* = 0.72). Figure 4b shows the mean voxel-level brain-predicted age for each participant against chronological age.

**Figure 4:**
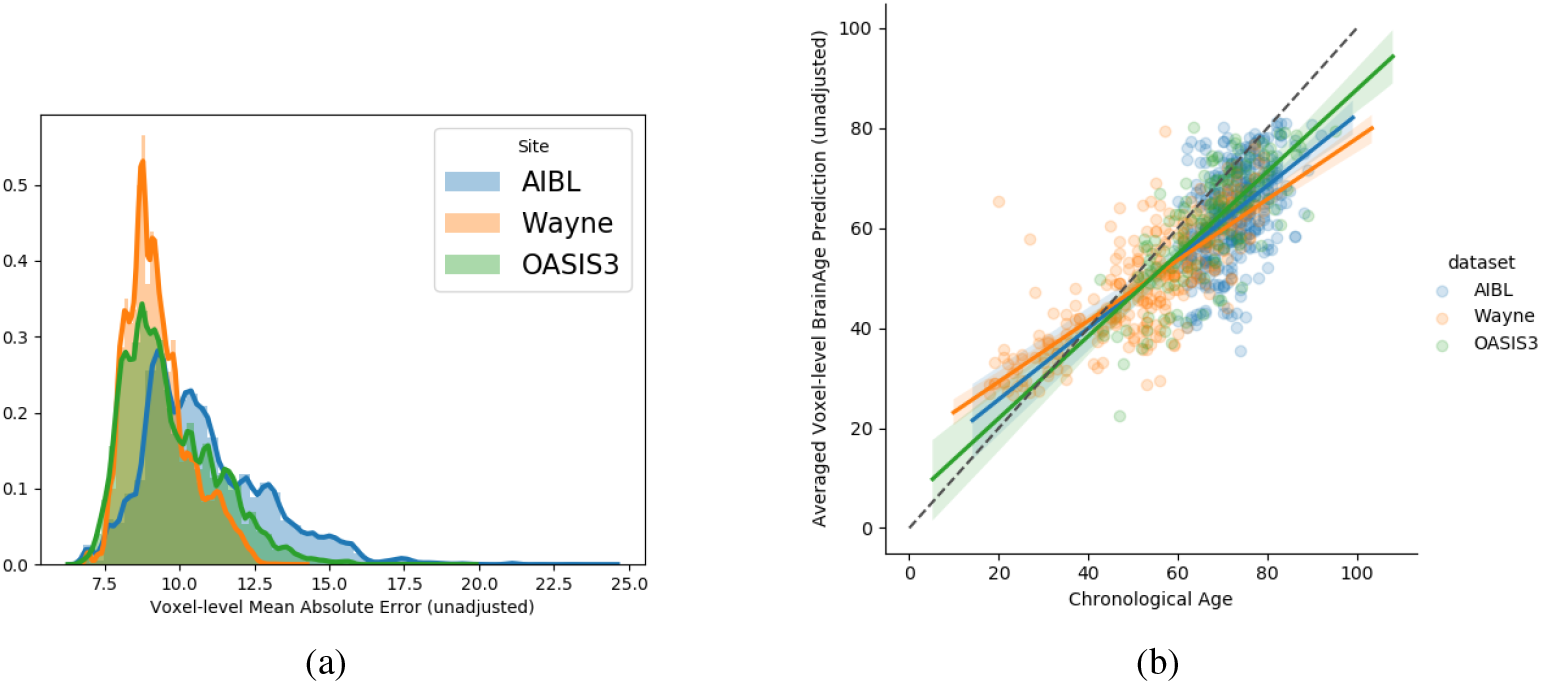
**a)** : Histogram of unadjusted, voxel-level MAE values across participants for each voxel. **b)**: global-level MAE (unadjusted) values plotted against chronological age.

**Figure 5:**
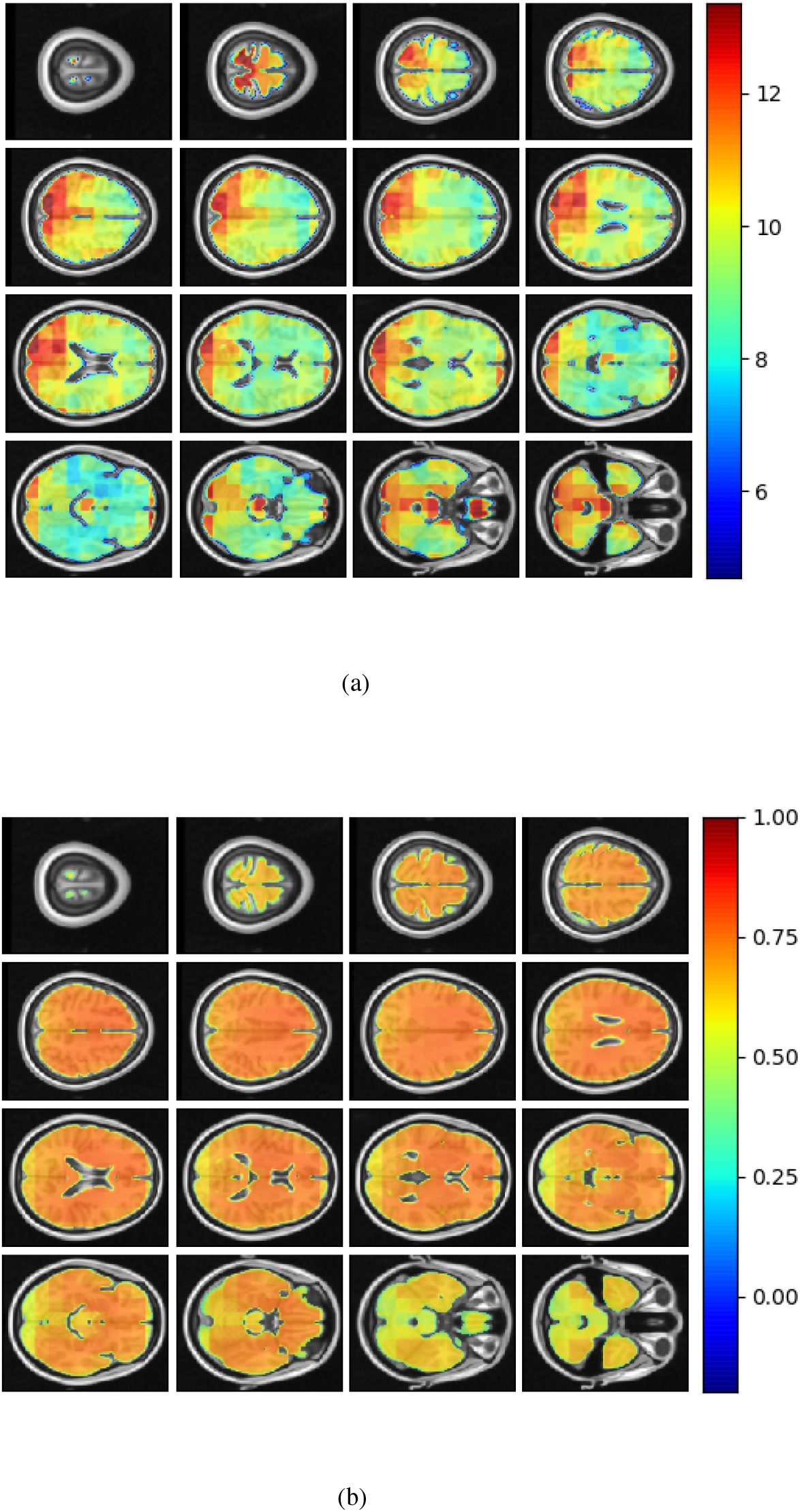
**a)**: Axial slices showing the spatial heterogeneity in unadjusted across participants voxel-level MAE values; **b)**: Pearson’s correlation coefficient between chronological age and voxel-level predicted brain-age (unadjusted) across participants.

### 3.2 Regional brain volumes and regional brain-PAD in healthy individuals

In this subsection we explored ROI-level results, based on the Harvard-Oxford atlas. We include cortical region results in the supplementary material (Table S2). From Table 3 we can observe that the amygdala, hippocampus and thalamus have the strongest negative Pearson’s correlation coefficients between ROI-level brain-PAD and ROI-level volumes (Figure 6).

**Table 3:** Pearson’s correlation coefficient (r) for different subcortical ROIs from the Harvard-Oxford atlas between ROI-level brain tissue volume and ROI-level brain-PAD.

| Subcortical ROI name | Left Hemisphere | Right Hemisphere |
| --- | --- | --- |
| Amygdala | -0.48 | -0.48 |
| Caudate | -0.13 | -0.19 |
| Hippocampus | -0.42 | -0.46 |
| Pallidum | -0.19 | -0.17 |
| Putamen | -0.32 | -0.32 |
| Thalamus | -0.34 | -0.43 |
| Accumbens | -0.23 | -0.30 |

**Figure 6:**
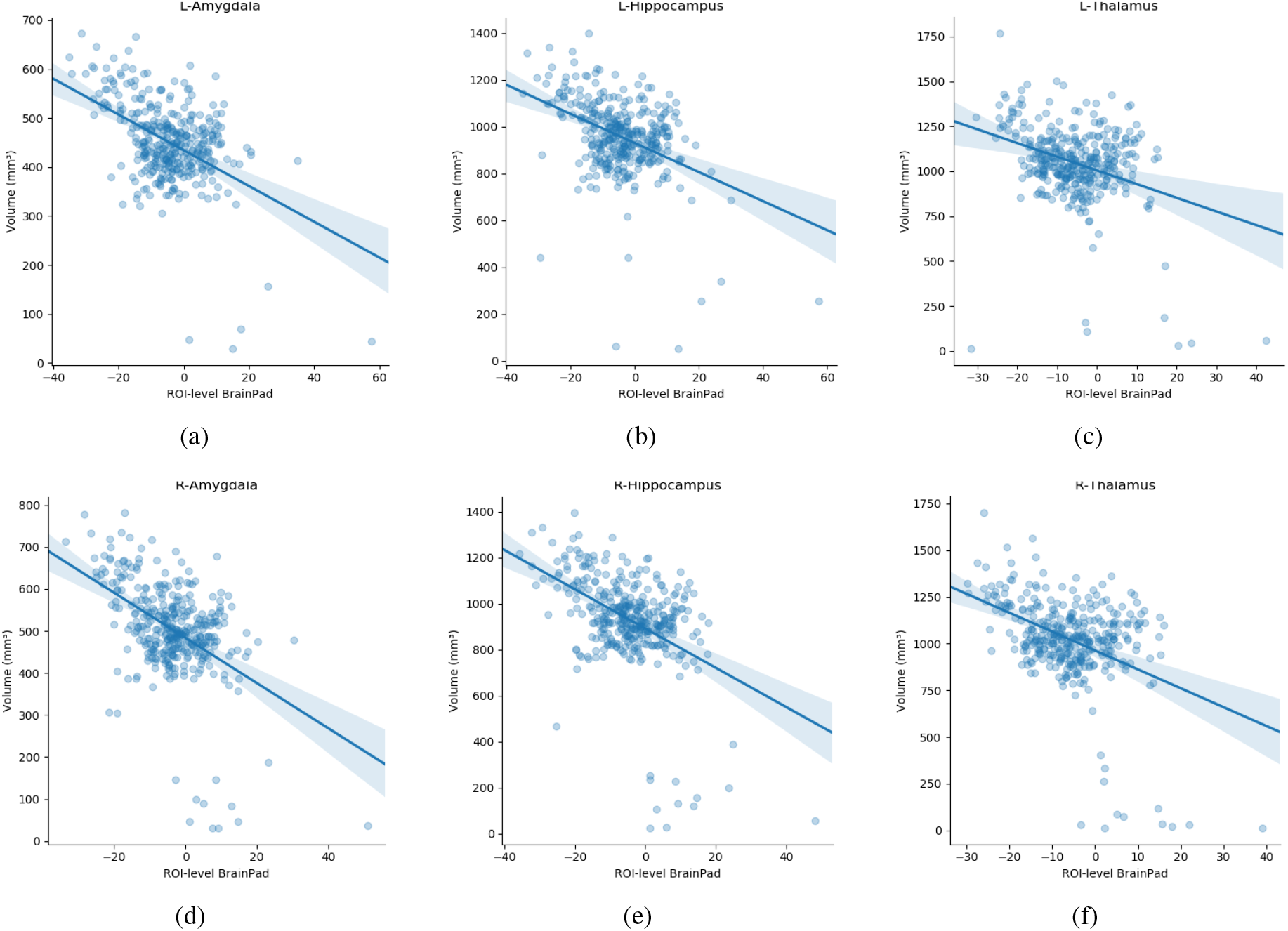
Scatterplots of ROI-level brain-PAD and brain volume (mm^3^). Volumes were generated using regional templates from the Harvard-Oxford atlas. **a)** Left amygdala, **b)** Left hippocampus. **c)** Left thalamus. **d)** Right amygdala. **e)** Right hippocampus. **f)** Right thalamus.

### 3.3 Reliability of local brain-age

Using voxel-level brain-age values for the Within-scanner (test-retest) and Scanner calibration (between-scanner) datasets, ICC was calculated per voxel. Test-retest reliability was very high with the vast majority of voxels having ICC<0.90 (median ICC = 0.98). This indicated very high reliability of local brain-age predictions within the same scanner. We observed comparatively lower ICC values at the extremities of the brain, see Figure 7b. This could be due to residual misregistration or partial volume effects. Between-scanner reliability was lower, with median voxel-level ICC = 0.62. Interestingly, the pattern of ICC varied across the brain, with higher values observed in the prefrontal cortex and lower values in more inferior regions, particularly the brainstem and cerebellum (Figure 7d).

**Figure 7:**
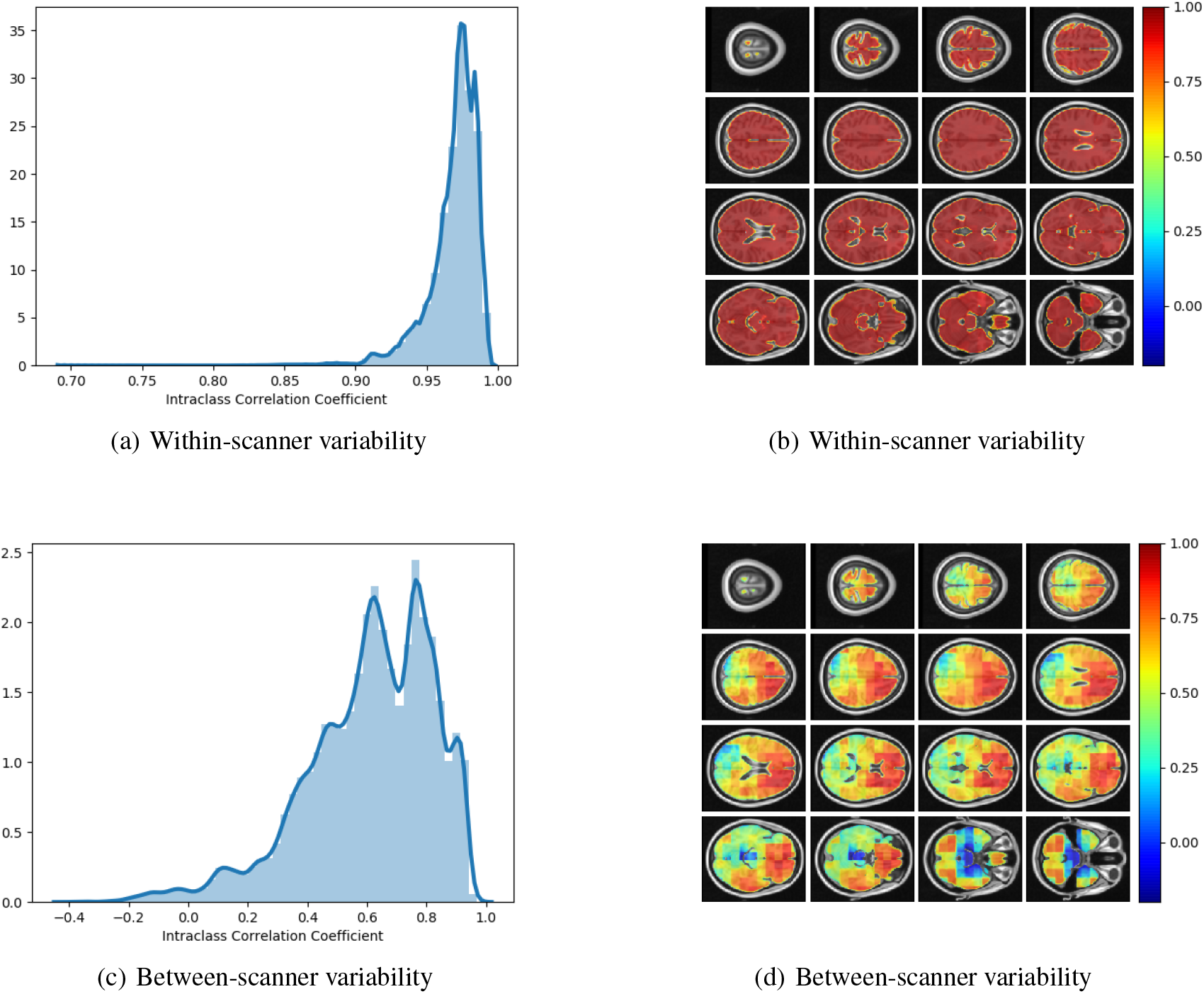
**a)**: Histogram of Intraclass Correlation Coefficients computed at voxel-level on STORM dataset. Values above 0.9 indicate strong agreements. **b)**: ICC values at different views on the axial plane on test-retest (i.e., within-scanner) dataset (n=20). **c)**: Histogram of Intraclass Correlation Coefficients computed at voxel-level on between-scanner reliability dataset (n=11, Siemens and Philips scanners). **d)**: ICC values at different axial slices from the between-scanner dataset.

### 3.4 Local brain-age differences between healthy controls, people with MCI and dementia patients

We examined patterns of local and global brain-age in people with MCI and dementia patients using cross-sectional data from OASIS3. Firstly, we investigated if the global-level (i.e., averaged within participant) brain-predicted age (adjusted) corresponds to previously reported differences from models that directly predict global brain age. We averaged voxel-level brain-age (adjusted) across voxels per individual to generate an adjusted global-level brain age and then calculate global-level brain-PAD. Global-level brain-PAD (adjusted) mean (± standard deviation) values were: −0.65 ± 7.46 (median=0.95) years for healthy controls, 3.07 ±4.29 (median = 2.83) years for stable MCI (sMCI), 5.77 ± 5.41 (median = 4.94) years for progressive MCI (pMCI) and 4.34 ± 6.78 (median = 4.63) years for AD patients.

We then assessed the significance of group differences using global-level brain-PAD values by performing independent two-sample Welch’s t-tests, finding significant differences between cognitively impaired groups and healthy controls in all cases (HC-AD *t* = −3.64, p = 0.0004, df = 88.96, Cohen’s *d* = −0.70; HC-sMCI *t* = −2.18, p = 0.0369, df = 29.29, Cohen’s *d* = −0.53; HC-pMCI *t* = −3.67, p = 0.0007, df = 38.66, Cohen’s *d* = −0.92). Comparisons between stable and progressive MCI patients and with AD patients were not significant: sMCI-pMCI p = 0.161, *t* = −1.44, df = 26.44, Cohen’s *d* = −0.54, AD-sMCI *t* = 0.83, p = 0.414, df = 20.64, Cohen’s *d* = 0.19, AD-pMCI *t* = −0.90, p = 0.3714, df = 28.51, Cohen’s *d* =-0.21.

Next, we examined local brain-PAD, summarising across all voxels within group. The mean voxel-level brain-PAD (adjusted) values were: healthy controls = −0.39 ± 0.85 (median = −0.44) years, sMCI = 3.07 ± 1.67(median = 3.266) years, pMCI = 5.45 ± 1.74 (median = 5.663) years for pMCI, AD patients = 4.01 ± 1.71 (median = 4.229) years (Figure 8b). We then compared groups based on these voxel-level brain-PAD values (adjusted) (Table 4 upper triangular part) using paired Welch’s t-test. Likewise, differences between participants MCI or dementia and healthy controls were significant (HC-sMCI *t* = −1284.67, p <0.0001, df = 723943.40, Cohen’s *d* = −2.60; HC-pMCI *t* = −2095.58, p < 0.0001, df = 707525.16, Cohen’s *d* = −4.25; HC-AD *t* = −1606.19, p <0.0001, df = 971564.04, Cohen’s *d* =-3.25). In contrast to the global-level results (lower triangle in Table 4), all pairwise differences between groups with MCI or dementia were significant (sMCI-pMCI *t* = −684.57, p <0.001, df = 970331.23, Cohen’s *d* = −1.38; sMCI-AD *t* = 272.44, p <0.001, df = 971564.04, Cohen’s *d* = 0.55; pMCI-AD *t* = −411.23, p <0.001, df = 971487.0, Cohen’s *d* = −0.83).

**Figure 8:**
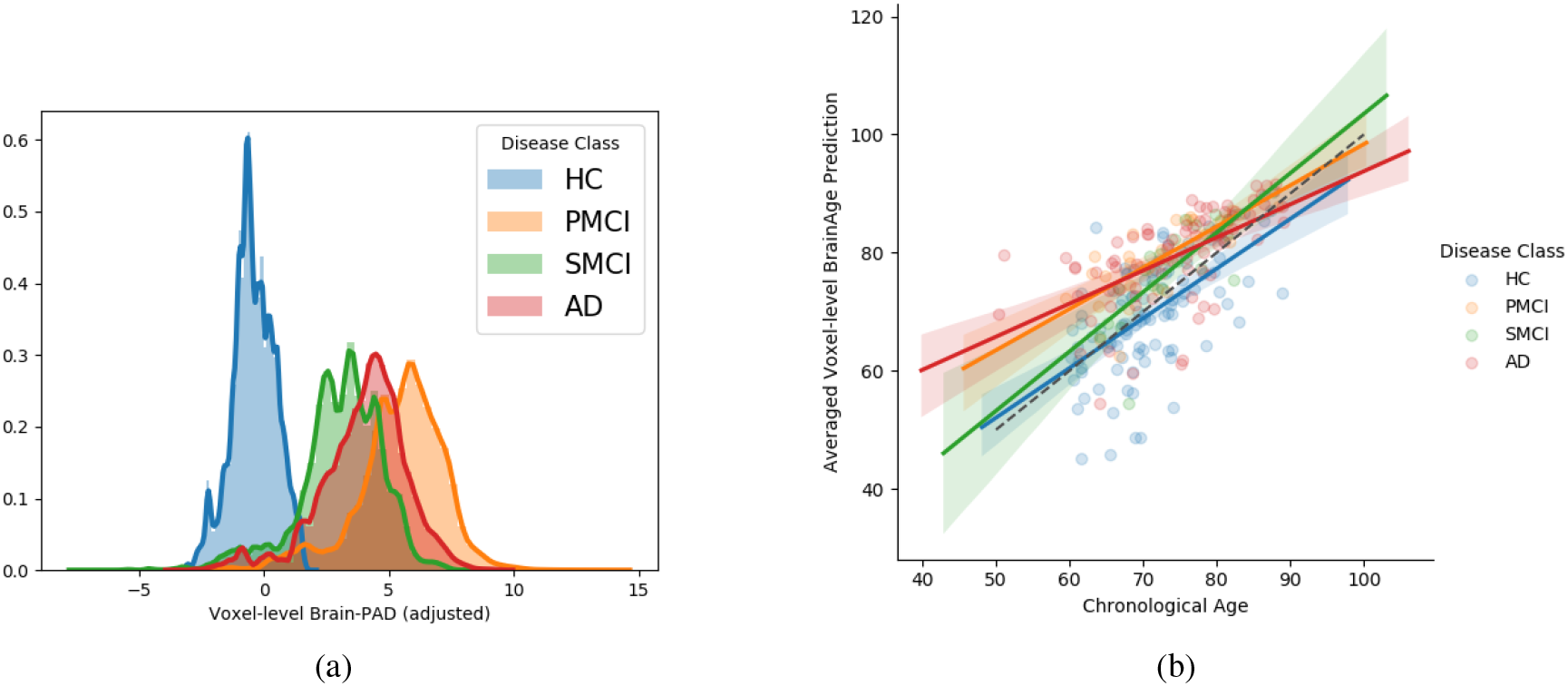
**a)** Histogram at voxel-level of brain-PAD scores of certain clinical groups from OASIS3. Brain-PAD after applying the bias-adjustment scheme is calculated for every voxel and then aggregated to the mean across all participants. Histograms in the plot are composed of the mean brain-PAD values for all voxels in the brain; **b)** Adjusted global-level predictions averaged across voxels for each participant; HC = Healthy controls, sMCI= stable MCI, pMCI = progressive MCI, AD = Alzheimer’s disease

**Table 4:** Group comparisons of brain-age in OASIS3 participants. Upper triangle: Voxel-level brain-age comparisons using paired Welch’s t-test results (t statistics value (p-value)) between disease groups. Lower triangle: Global-level brain-age comparisons using independent Welch’s t-test results (t statistics value (p-value)) between disease groups.

| Disease Groups | HC | sMCI | pMCI | AD |
| --- | --- | --- | --- | --- |
| HC | - | -1284.67<br>(<0.001) | -2095.58<br>(<0.001) | -1606.19<br>(<0.001) |
| sMCI | -2.18<br>(0.0369) | - | -684.57<br>(<0.001) | 272.44<br>(<0.001) |
| pMCI | -3.67<br>(0.0007) | -1.44<br>(0.1611) | - | -411.23<br>(<0.001) |
| AD | -3.64<br>(0.0004) | 0.83<br>(0.4141) | -0.90<br>(0.3714) | - |

From Figure 9 we can observe that local brain-age model is able to detect group differences across the whole brain when comparing healthy controls with AD patients or comparing the pMCI group with the sMCI group (after correction for multiple comparisons). Other group contrasts showed more varied spatial patterns of significant voxels. From Figure 10 we can observe that the largest differences are in the temporal lobe and subcortical regions when comparing AD patients to healthy controls. For a more in-depth look at differences between disease groups, we extended the analysis to investigate atlas-based subcortical ROIs. The nucleus accumbens, putamen, pallidum and hippocampus were the most discriminative ROIs in terms of Cohen’s *d* scores both for separating AD patients from healthy controls and stable from progressive MCI (Table 5). We also include histograms of the local brain-PAD scores for each disease group per subcortical ROI to visualise the different distributions that drive the report effect sizes (Figure 11). For example, the high Cohen’s *d* values for the nucleus accumbens may be due to the low variance in brain-PAD values in this small region. We have provided similar graphics for the cortical regions in the supplementary material (Figure S6).

**Figure 9:**
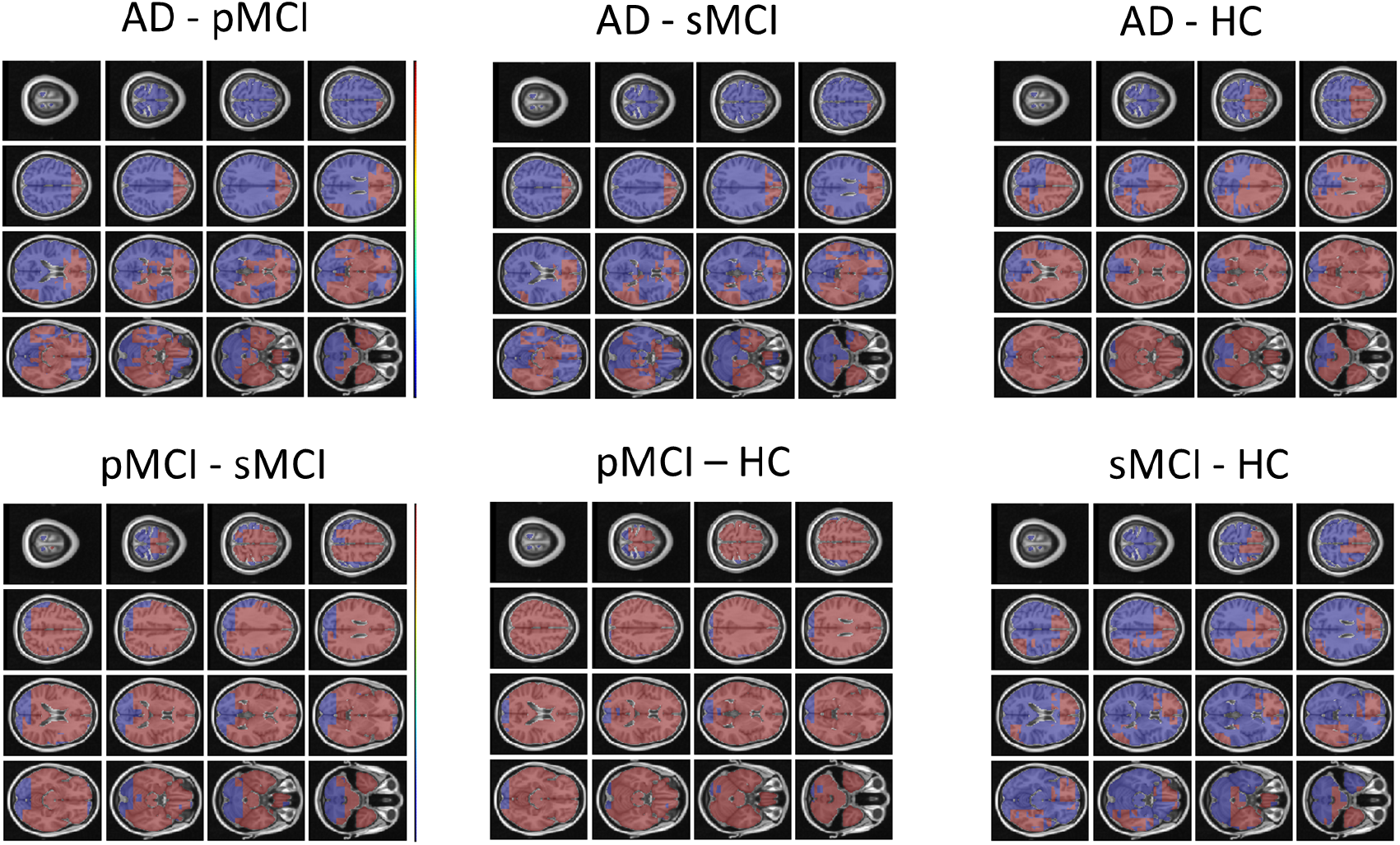
FSL Randomise maps for different combinations of clinical groups in cross-sectional OASIS3. Red colored voxels indicate a significant statistical t-test after correcting for multiple comparisons. Blue regions were not significant after correction. HC=Healthy Controls; pMCI = progressive MCI; sMCI = stable MCI; AD=Alzheimer’s Disease.

**Figure 10:**
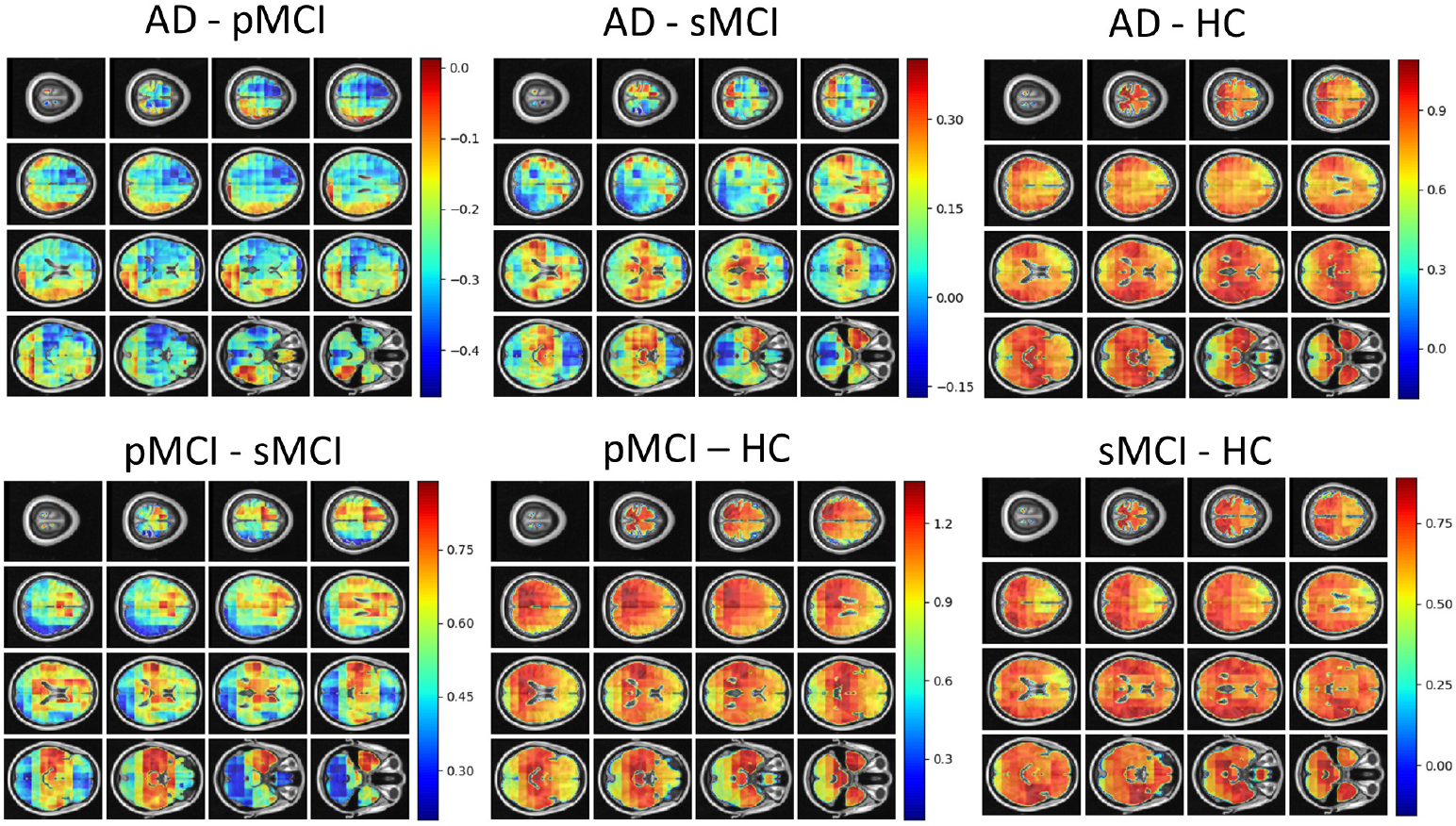
Cohen’s *d* maps for different combinations of cross-sectional comparisons of clinical groups in OASIS3. Positive values indicate a positive effect for the first group. HC=Healthy Controls; pMCI = progressive MCI; sMCI = stable MCI; AD=Alzheimer’s Disease.

**Table 5:**
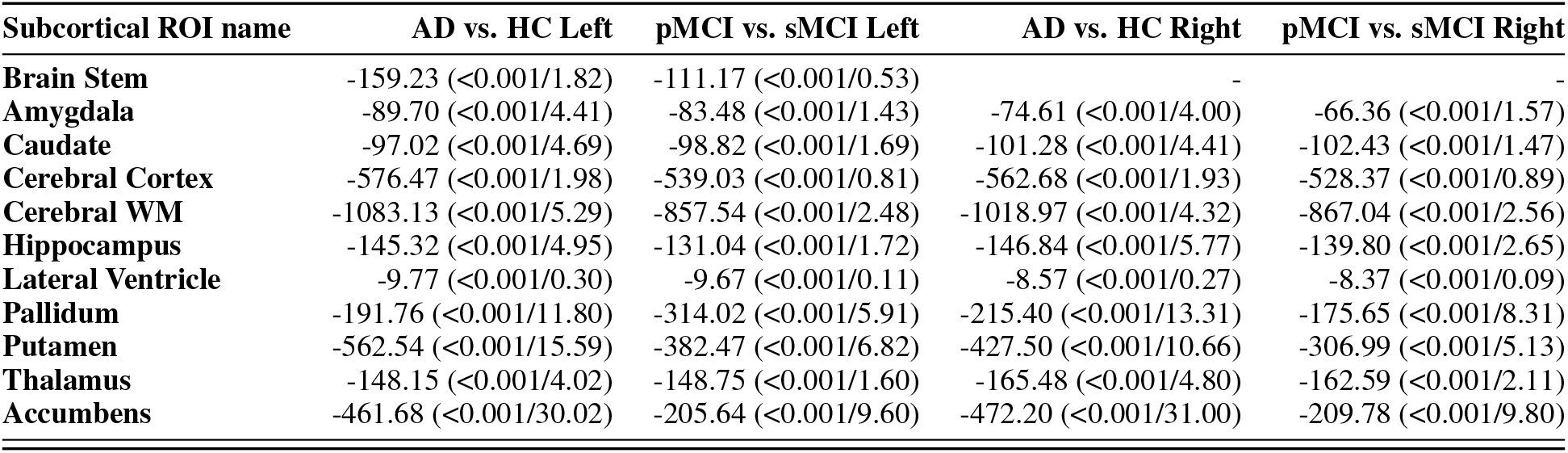
Welch’s t-test statistic (p-value/Cohen’s *d*) values for different subcortical ROIs from the Harvard-Oxford atlas. For Cohen’s *d*, higher values indicate a positive effect size for the first disease group specified.

**Figure 11:**
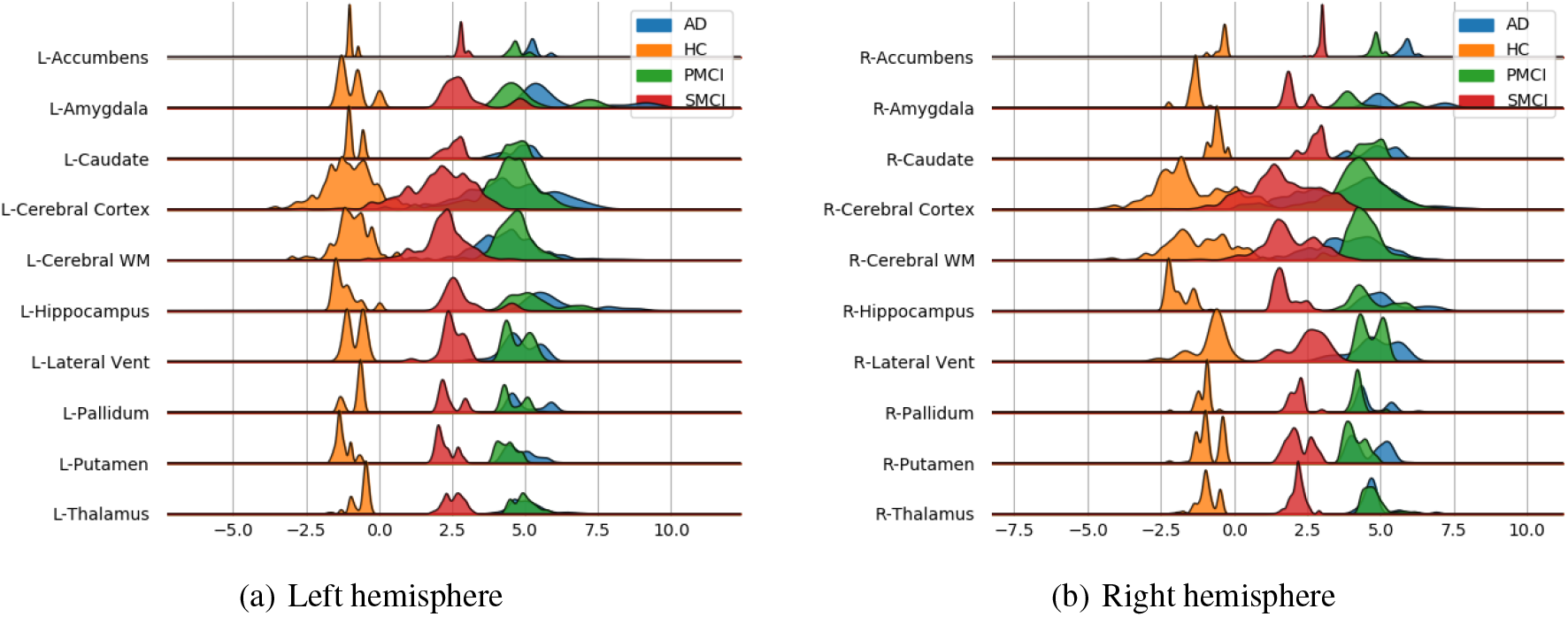
Subcortical ROI-based difference in voxel level brain-PAD scores averaged across participants from clinical groups from OASIS3. X axis shows brain-PAD values within the given ROI; HC = Healthy controls, sMCI= stable MCI, pMCI = progressive MCI, AD = Alzheimer’s disease

## 4 Discussion

In this paper, we introduced a novel deep-learning framework capable of reliably predicting age from neuroimaging data at a local neuroanatomical level. Training the powerful U-net architecture on n=3463 healthy people, we present the first proof-of-concept, to our knowledge, that generating such localised brain-age predictions is feasible. While average performance of our model (MAE = 9.94 ± 1.73 years) is below what has been reported with purely global (MAE 3 years, [40]), slice-level (MAE between 5-7.5 years, [41]), or patch-level (MAE 2.5-4 years, [23]), we show both high reliability and reasonable generalisability to three entirely independent datasets (OASIS3, AIBL, Wayne State). Importantly, we achieved a resolution of 23^3^ voxels, substantially more fine-grain than previous patch-level work (64^3^ voxels [23]). In fact, we were able to generate voxel-level prediction, though as within-block homogeneity was high, our effective resolution was lower than single voxel. Future improvements to network architecture may be able to improve the effective resolution still further.

Even though the mean global performance of the local brain-age model was relatively poor, the model still demonstrated sensitivity to cognitive impairment and dementia, suggesting that despite the noise at test time, the relevant signal can still be observed. Previous work involving brain-age and dementia have obtained “brain-AGE” scores of −0.2 years for sMCI, 6.2 years for pMCI and 6.7 years for AD on the ADNI dataset [10]. Our results are generally in line with these previous findings, though here, we observed “older-appearing” brains our sMCI group (mean brain-PAD = 2.8 years). Moreover, we were able to generate spatial maps of brain-PAD for each individual, showing how the patterns of brain-ageing may vary across the brain in a single patient. At the local level, we observed widespread patterns of group differences in brain-PAD maps, including when comparing sMCI and pMCI groups, suggesting that brain-ageing is more pronounced in those MCI patients who go on to develop dementia within three years.

It has been commonly reported that the early stages of AD involve atrophy in the medial temporal lobe (MTL) including the hippocampus, and the amygdala, entorhinal and parahippocampal cortices [42, 43, 44, 45]. Our voxel-level analysis showed brain-PAD differences between healthy controls and people with MCI in these key AD-related regions. Furthermore, the ROI-level analysis of brain-PAD should widespread differences with particularly strong effects in the nucleus accumbens, putamen, pallidum, hippocampus and amygdala as well as cortical and ventricular regions. While further research is required to further improve the model performance and spatial precision, these results suggest that the local brain-age predictions are sensitivity to local patterns of brain atrophy.

The validity of the predictions from the local brain-age model are further supported by the observed significant negative correlations between ROI volumes and ROI-level brain-PAD. In a similar analysis, Levakov et al. [19] identified the lateral ventricles, inferior lateral ventricles, 3rd ventricles, non-ventricles CSF and left/right choroid plexus as the ROIs (using the FreeSurfer Desikan-Killiany atlas) having the strongest relationships between age normalised volume and brain-age “gap”. Here, we also show relationships in GM ROIs (e.g., amygdala, hippocampus, thalamus, parahippocampal gyrus (anterior division), inferior temporal gyrus, temporal-occipital part, intracalcarine cortex). As lower brain volumes are associated with ageing, the observed negative relationships between ROI volume and ROI brain-PAD suggests that indeed the ROI-level brain-PAD captures some age-related variance.

As biomarker of brain health, brain-age models may have clinical utility, either prognostically or in the context of clinical trials of neuroprotective treatments. While previous studies have reported standardised effect sizes from global brain-age, we used atlas ROIs to summarise regional values of local brain-PAD and generated Cohen’s *d* values from pairwise group comparisons. Using conventional hippocampal volumetric measures, Henneman et al. [46] reported baseline effect size of 0.73 when comparing controls and MCI groups, and 0.33 when comparing people with MCI and AD patients. With our local brain-age framework, the control-MCI effect size for the hippocampus (average bilaterally) was *d* = 5.45 and the MCI-AD effect size was *d* = 0.48. Using voxel-based morphometry, [47] generated Cohen’s *d* values for the hippocampus (d = 0.6) and amygdala (d = 0.45), when comparing stable and progressive MCI patients. Here, our local brain-age framework resulted in *d* = 2.18 for the hippocampus and *d* =1.5 for the bilateral amygdala in the same context. This suggests that use of the brain-age paradigm to capture local age-related changes, relative to a healthy ageing model, could increase statistical power in experimental research and clinical trials, relative to conventional volumetric imaging biomarkers. Potentially, the ROI-based brain-PAD values could even be used in a classification framework to distinguish between people with stable or progressive MCI.

Out proposed U-Net local brain-age framework has some strengths and weaknesses. Our model was assessed on a large multi-site testing set with a flat distribution of chronological age across the adult lifespan (18-90 years; Figure S1), a wider interval than a number of studies that rely on UK Biobank [40, 23, 25] or other narrower-age range studies. Our model showed excellent test-retest reliability, giving confidence that the model could be applied longitudinally to assess individual patterns of brain-ageing changes. However, the between-scanner reliability was moderate, similar to our previous work using deep learning to predict brain age [33]. In the latter work, brain-age prediction was performed directly on raw MRI scans, hence the deep learning model may be overfitting to some site or effects. One might expect that image pre-processing may partially ameliorate these site effects. However, this is not uniformly the case, as previous research has demonstrated [48]. Consequently, one drawback of the current algorithm is the requirement to have a healthy population from a given clinical site to use as a control group, as site or scanner effects may result in the local brain-PAD distribution not being centred at zero. In Supplementary Figures S7 and S8 we show that these scanner effects have only a marginal effect on the statistical comparisons between MCI or dementia groups using the AIBL dataset in reference to our main results from OASIS3. Nevertheless, harmonisation of scanner and site is a key direction for future work as the removal of residual scanner effects is likely to improve model generalisability considerably and is an important prerequisite for the clinical adoption of neuroimaging biomarker pipelines.

We trained our regression U-Net with the ground truth objective at a voxel-level (given by a three-dimensional block filled with the chronological age), in order to encourage the network to emphasise the context encoded in its lower layers. As the individual voxel location we are aiming to obtain a prediction for is not necessarily related to the imposed ground truth output, the U-Net architecture is biased towards using the context information. Hence, in the worst case scenario where no voxel-level relationship is learned, the true resolution of our voxel-level predictions is actually blocks of 23^3^ voxels. The final output field-of-view (FOV) was calculated starting from the first convolutional layer where the FOV is 3^3^ voxels, which gets increased by 2^3^ voxels per convolutional operation in the downstream part of the U-Net. The average pooling layers increase the FOV by 1^3^ voxels, since their stride is set to 1, while the upsampling layers do not increase the FOV as they merely repeat existing information. Lastly, the first convolutional layer in the upstream layers only adds 1^3^ voxels (since a 2 * 2 * 2 block inside the operating field of the 3 * 3 * 3 filter contains the same repeated information, hence no increase in the FOV) whereas the second adds 3^3^ voxels (since a 3 * 3 * 3 filter will have access to 3 additional voxels stemming from the upsampling layer). While this means that our resolution is not necessarily at the voxel-level, 23^3^ voxels is still substantially higher resolution compared to existing models in literature. In the 3D block approach of Bintsi et al. [23], blocks are much larger, 64^3^ voxels voxels. Hence, any block-level age prediction will be biased towards the global-level brain age prediction as the blocks include a substantial portion of the overall brain. Moreover, in splitting the whole brain into blocks, naturally some blocks will include non-brain tissue or empty space, which will naturally reduce the amount of discriminative information present there, reducing the validity of results for regions within the respective block.

We demonstrated how our U-net framework can predict age from neuroimaging data. However, this approach could be trained on any continuous or categorical outcome measure to generate individual maps of how given outcomes vary across the brain. For example, one could generate spatial maps of predicted values of fluid biomarkers (e.g., amyloid, tau), genotype or polygenic risk score, cognitive measures (e.g., MMSE scores). Such an approach could be used as an alternative to techniques like VBM, to provide mechanistic insights into the relationship between local brain regions and individual deviations from healthy/normal levels of a given outcome measure. While VBM is the *de facto* method to quantitatively assess differences between groups at voxel-level [49], we believe the local brain-age framework is complementary to this. In VBM analysis, one assesses the statistical models at a voxel level based on volume or intensity, though local context is only really accounted for at the cluster inference stage. Brain-PAD implicitly measures this deviation of the diseases group from what constitutes a normative pattern of ageing, by placing the participant on a distribution of normative ageing for a given local area (i.e., the voxel and its local context). We leave for further work the comparison between VBM and local brain-age.

One potential direction to take local brain-age further is in disease subtyping. Local brain-PAD maps could be used as input to clustering algorithms with the goal of identifying subgroups of patients that have spatially similar patterns of brain ageing. The putative subgroups may undergoing distinct pathological processes that effect different regions of the brain and may have different trajectories of disease progression or may respond differently to treatments. Such approaches have been applied to volumetric brain maps before [50], but the addition of brain-PAD information as a local index of age-adjusted brain health could increase sensitivity, as has been seen in global brain-age research [51].

### Conclusion

We have introduced a new deep learning framework that is capable of reliably estimating brain-age with high spatial resolution, providing information on spatial patterns of age-related changes to brain volume. We were able to demonstrate the potential clinical relevance of the model by mapping differences in local and regional brain-PAD scores in patients with cognitive impairment and dementia. This work illustrates how the sensitivity of conventional global brain-age analysis can be augmented with individualised spatial maps offering potential mechanistic insights, with the goal of opening the “black box” of the machine learning algorithms that underpin the brain-age paradigm.

## Data and code availability statement

The data used in these experiments are available on application to the relevant studies. The code used is available at https://github.com/SebastianPopescu/U-NET-for-LocalBrainAge-prediction alongside the pre-trained models.

## 5 Acknowledgments

SGP is funded by an EPSRC Centre for Doctoral Training studentship award to Imperial College London. BG received funding from the European Research Council (ERC) under the European Union’s Horizon 2020 research and innovation programme (grant agreement No 757173, project MIRA, ERC-2017-STG). DJS is supported by the NIHR Biomedical Research Centre at Imperial College Healthcare NHS Trust and the UK Dementia Research Institute (DRI) Care Research and Technology Centre. JHC acknowledges funding from UKRI/MRC Innovation Fellowship (MR/R024790/2).

## 6 Disclosure of competing interests

BG has received grants from European Commission and UK Research and Innovation Engineering and Physical Sciences Research Council, during the conduct of this study; and is Scientific Advisor for Kheiron Medical Technologies, Advisor and Scientific Lead of the HeartFlow-Imperial Research Team, and Visiting Researcher at Microsoft Research. JC is a shareholder in and Scientific Advisor to BrainKey and Claritas Healthcare, both medical image analysis software companies.

## 7 Credit authorship contribution statement

Sebastian G. Popescu: Conceptualisation, Methodology, Software, Formal analysis, Investigation, Writing - original draft, Writing - review & editing, Visualisation. Ben Glocker: Conceptualisation, Methodology, Writing - review & editing, Supervision. David J.Sharp: Conceptualisation, Methodology, Writing - review & editing, Supervision. James H. Cole : Conceptualisation, Methodology, Writing - review & editing, Supervision.

## Notes

### Competing Interest Statement

The authors have declared no competing interest.

### Summary of Updates

Additional results and clarifications.

