## supplementary material for "Local brain-age: A U-Net model"

#### A Brain-Age Healthy Control (BAHC) dataset

| Cohort | N | Age mean (SD) | Age range | Sex M/F | Repository details | Scanner (Field strength) | Scan | Voxel dimensions |
|---|---|---|---|---|---|---|---|---|
| ABIDE | 184 | 25.93 (6.66) | 18-48 | 161/23 | INDI | Various (all 3T) | MPRAGE | Various |
| Beijing Normal University | 179 | 21.25 (1.92) | 18-28 | 72/107 | INDI | Siemens (3T) | MPRAGE | 1.33x1.0x1.0 |
| Berlin School of Brain Mind | 49 | 30.99 (7.08) | 20-60 | 24/25 | INDI | Siemens Tim Trio (3T) | MPRAGE | 1.0x1.0x1.0 |
| CADDementia | 12 | 62.33 (6.26) | 55-79 | 9/3 | link | GE Signa (3T); 3D IR-FSPGR | 0.9x0.9x1.0 | |
| Cleveland Clinic | 31 | 43.55 (11.14) | 24-60 | 11/20 | INDI | Siemens Tim Trio (3T) | MPRAGE | 2.0x1.0x1.2 |
| ICBM | 322 | 24.84 (5.14) | 24-60 | 177/145 | LONI IDA | Siemens Magnetom (1.5T) | MPRAGE | 1.0x1.0x1.0 |
| IXI | 561 | 48.62 (16.49) | 20-86 | 250/311 | link | Philips Intera (3T); Philips Gyroscan Intera (1.5T); GE Signa (1.5T) | T1-FFE; MPRAGE | 0.9375x0.93751x1.2 |
| MCIC | 93 | 32.49 (11.95) | 18-60 | 64/29 | COINS | Siemens Sonata/Trio (1.5/3T); GE Signa (1.5T) | MPRAGE; SPGR | 0.625x0.625x1.5 |
| MIRIAD | 23 | 69.66 (7.18) | 58-85 | 12/11 | link | GE Signa (1.5T) | 3D IR-FSPGR | 0.9375x0.93751x1.5 |
| NEO2012 (Adelstein, 2011) | 39 | 29.59 (8.38) | 20-49 | 18/21 | INDI | Siemens Allegra (3T) | MPRAGE | 1.0x1.0x1.0 |
| Nathan Kline Institute (NKI) / Rockland | 160 | 41.49 (18.08) | 18-85 | 96/64 | INDI | Siemens Tim Trio (3T) | MPRAGE | 1.0x1.0x1.0 |
| OASIS | 288 | 44.06 (23.04) | 18-90 | 106/188 | link | Siemens Vision (1.5T)* | MPRAGE | 1.0x1.0x1.25 |
| WUS (Power, 2012) | 24 | 23.04 (1.42) | 20-24 | 4/20 | INDI | Siemens Tim Trio (3T) | MPRAGE | 1.0x1.0x1.0 |
| TRAIN-39 | 36 | 22.67 (2.56) | 18-28 | 11/25 | INDI | Siemens Allegra (3T) | MPRAGE | 1.33x1.33x1.3 |
| Training set total | 2001 | 36.95 (18.12) | 18-90 | 1016/985 | - | - | - | - |

Table S1: ABIDE = Autism Brain Imaging Data Exchange; ICBM = International Consortium for Brain Mapping; IXI = Information eXtraction from Images; MCIC = MIND Clinical Imaging Consortium; MIRIAD = Minimal Interval Resonance Imaging in Alzheimer's Disease; OASIS = Open Access Series of Imaging Studies; INDI = International Neuroimaging Data-sharing Initiative (http://fcon_1000.projects.nitrc.org); COINS = Collaborative Informatics and Neuroimaging Suite (http://coins.mrn.org); LONI = Laboratory of Neuro Imaging Image Data Archive (https://ida.loni.usc.edu); ABIDE consortiums comprising data from various sites with different scanners/parameters; *OASIS scans were acquired four times and then averaged to increase signal-to-noise ratio.

#### B Chronological age histogram for datasets

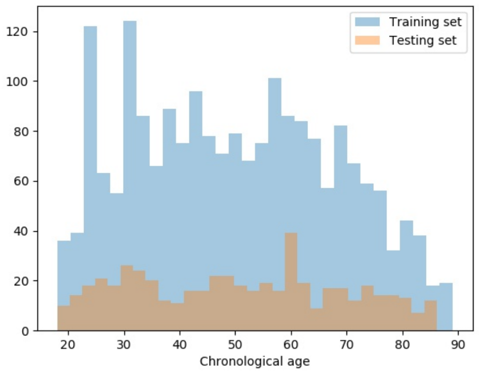


(a) Healthy datasets

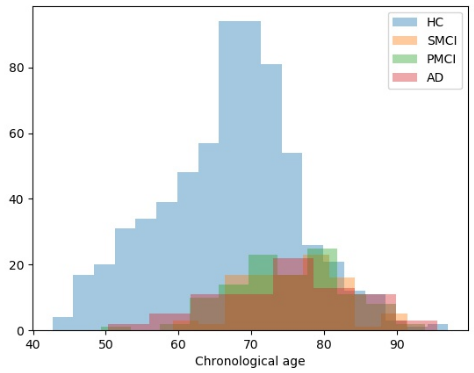


(b) OASIS3

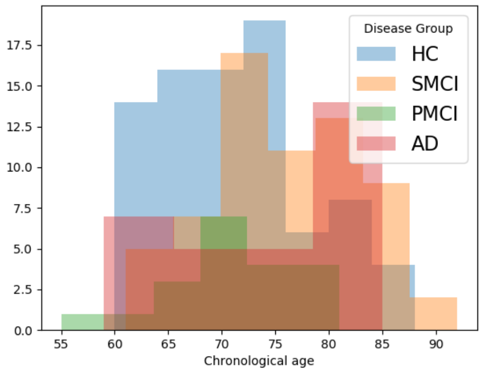


(c) AIBL

Figure S1: Chronological age histograms for healthy and clinical datasets. Healthy subjects datasets pooled were constructed such as to have a relatively flat training and testing chronological age histogram. Subjects included have chronological age varying from 18 to 90 years old, thereby our results are able to catch a clearer picture of our model's effectiveness across the whole aging spectrum.

#### C Cortical level relationship between volumetric mass and regional BrainPAD

| **Cortical ROI name** | **Pearson's *r*** |
|---|---|
| Frontal Pole | -0.24 |
| Insular Cortex | -0.31 |
| Superior Frontal Gyrus | -0.31 |
| Middle Frontal Gyrus | -0.29 |
| Inferior Frontal Gyrus, pars triangularis | -0.37 |
| Inferior Frontal Gyrus, pars opercularis | -0.37 |
| Precentral Gyrus | -0.25 |
| Temporal Pole | -0.33 |
| Superior Temporal Gyrus, anterior division | -0.37 |
| Superior Temporal Gyrus, posterior division | -0.31 |
| Middle Temporal Gyrus, anterior division | -0.40 |
| Middle Temporal Gyrus, posterior division | -0.37 |
| Middle Temporal Gyrus, temporooccipital part | -0.37 |
| Inferior Temporal Gyrus, anterior division | -0.37 |
| Inferior Temporal Gyrus, posterior division | -0.39 |
| Inferior Temporal Gyrus, temporooccipital part | -0.43 |
| Postcentral Gyrus | - 0.29 |
| Superior Parietal Lobule | -0.31 |
| Supramarginal Gyrus, anterior division | -0.32 |
| Supramarginal Gyrus, posterior division | -0.30 |
| Angular Gyrus | -0.32 |
| Lateral Occipital Cortex, superior division | -0.26 |
| Lateral Occipital Cortex, inferior division | -0.34 |
| Intracalcarine Cortex | -0.42 |
| Frontal Medial Cortex | -0.38 |
| Juxtapositional Lobule Cortex | -0.41 |
| Subcallosal Cortex | -0.36 |
| Paracingulate Gyrus | -0.31 |
| Cingulate Gyrus, anterior division | -0.26 |
| Cingulate Gyrus, posterior division | -0.34 |
| Precuneous Cortex | -0.34 |
| Cuneal Cortex | -0.42 |
| Frontal Orbital Cortex | -0.30 |
| Parahippocampal Gyrus, anterior division | -0.50 |
| Parahippocampal Gyrus, posterior division | -0.31 |
| Lingual Gyrus | -0.37 |
| Temporal Fusiform Cortex, anterior division | -0.41 |
| Temporal Fusiform Cortex, posterior division | -0.39 |
| Temporal Occipital Fusiform Cortex | -0.32 |
| Occipital Fusiform Gyrus | -0.40 |
| Frontal Operculum Cortex | -0.25 |
| Central Opercular Cortex | -0.31 |
| Parietal Operculum Cortex | -0.27 |
| Planum Polare | -0.31 |
| Heschls Gyrus | -0.32 |
| Planum Temporale | -0.33 |
| Supracalcarine Cortex | -0.41 |
| Occipital Pole | -0.16 |

Table S2: Pearson's correlation coefficient for different cortical ROIs from the Oxford-Harvard atlas between ROI-level brain tissue volume and ROI-level brain-PAD.

In this subsection we explored cortical ROI-level results based on the Harvard-Oxford atlas. From Table S2 we can observe that the Parahippocampal Gyrus (anterior division), Inferior Temporal Gyrus (temporooccipital part), Intracalcarine Cortex and Cuneal Cortex have the strongest negative Pearson's correlation coefficients between ROI-level brain-PAD and ROI-level volumes. Compared to [1],where they obtained absolute Pearson's correlation coefficients less than 0.15 for Cortical GM, our method is consistently able to obtain absolute values higher than 0.15, with the lowest one being achieved in the Occipital Pole (-0.16).

### D Mass univariate linear regression bias adjustment framework

As pointed out in [2], most studies in brain age prediction that intend to eliminate age related bias for further group comparison utilize a variant relating on linear regression with covariate given by the brain predicted age for the respective subject. We have also attempted this pathway for this project albeit unsuccessfully. We reiterate that after averaging the brain age delta across voxels for subjects, the brain age delta has the negative pattern as observed in [3], however upon looking at voxel level, this negative pattern gets slightly lost due to high variance or some outlier points which influence the linear regression fit (Figure S2). Potential avenue for further research is to instead use a non-parametric model that more easily ignores outlier points such as generalized additive models or Gaussian Processes.

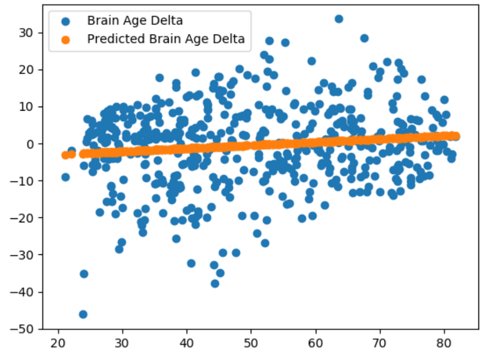


(a)

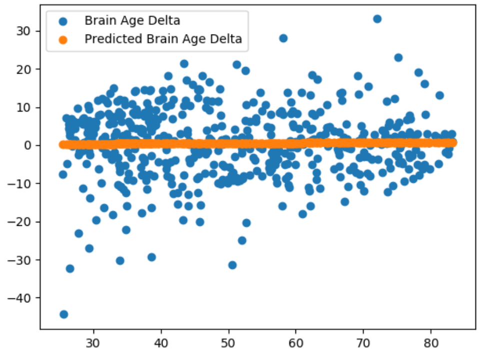


(b)

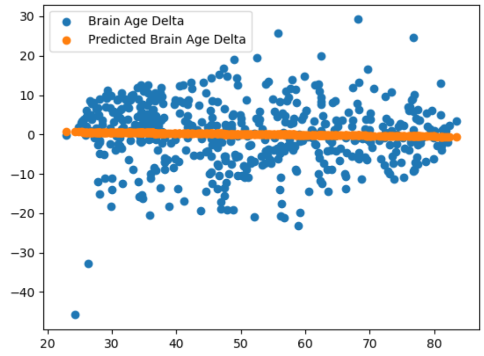


(c)

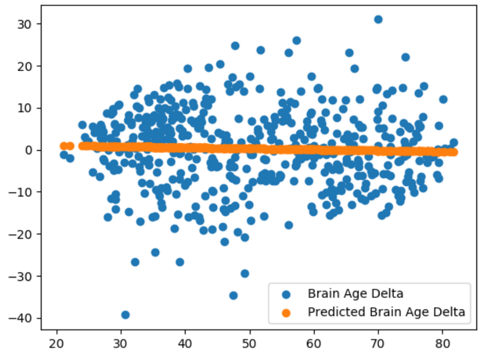


(d)

Figure S2: Scatter plots of chronological age and brain age delta. Results from randomly sampled voxels in the brain. With orange we plot the fitted linear regression used for bias adjustment.

### E Exploring bias-adjustment effectiveness at voxel-level

From Figure S3 we notice that our proposed bias-adjustment method has managed to eliminate the age-related bias in predictions, irrespective of dataset origin. For Wayne State, our bias adjustment framework increased the Pearson correlation (between predicted age and chronological age) from 0.61 to 0.76, for OASIS3 from 0.53 to 0.63, respectively for AIBL from 0.23 to 0.35. Moreover, in terms of actual precision of local brain-age, we obtain a distribution centered around 0 years mean absolute error before bias-adjustment. What is more important for clinical purposes is that the voxel-level brain-PAD scores for healthy controls are centred around 0, which indicates that our algorithm is capable to detect normative samples. Moreover, from Figure S4 we can notice that age-related bias is almost completely eliminated in 50 randomly picked voxels for subjects between 60 and 90 years old, which further shows the reliability of statistical inference performed on subgroups within that age range.

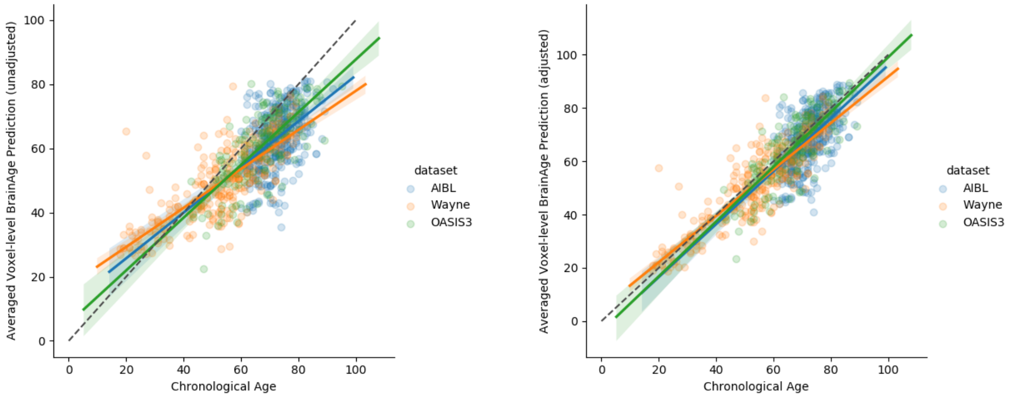


(a) (b)

Figure S3: **Left**: global-level MAE (unadjusted) values plotted against chronological age. **Right**: global-level MAE (adjusted) values plotted against chronological age.

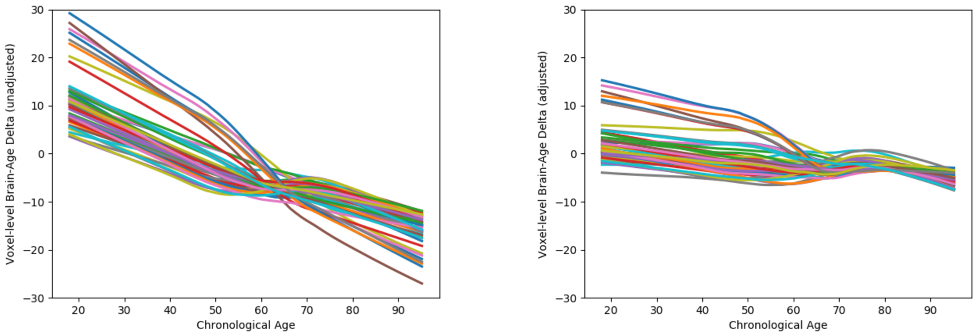


(a) (b)

Figure S4: LOWESS estimates of brain-age delta for 50 randomly sampled voxels in the brain before/after bias adjustment.

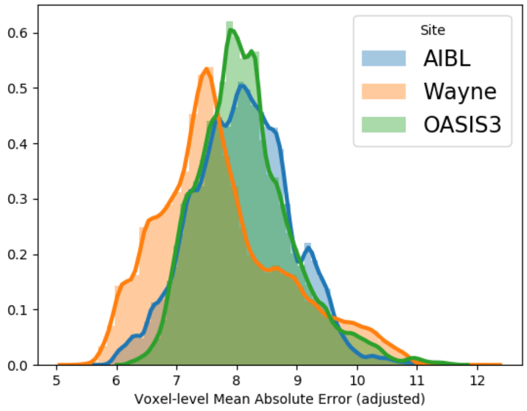


Figure S5: Per voxel averaged across subjects Mean Absolute Error (adjusted)

#### F Regional Cortical level differences between varying degrees of cognitive impairment

In this subsection our aim is to gain a deeper understanding of the spatial specificity of local brain-age by inspecting the differences in age-related brain atrophy at a cortical ROI level. The Parietal Operculum Cortex, Heschls Gyrus, Cingulate Gyrus (anterior division) and Central Opercular Cortex were the most discriminative ROIs in terms of Cohen's *d* scores for both separating AD patients from healthy controls, respectively stable from progressive MCI (Table S3 and Figure S6).

| **Cortical ROI name** | **AD vs. HC** | **pMCI vs sMCI** |
|---|---|---|
| **Frontal Pole** | -287.80 («0.001/1.70) | -261.39 («0.001/0.55) |
| **Inf-Frontal Gyrus, pars opercularis** | -143.08 («0.001/3.22) | -144.39 («0.001/1.31) |
| **Inf-Frontal Gyrus, pars triangularis** | -120.09 («0.001/2.58) | -120.39 («0.001/0.99) |
| **Insular Cortex** | -423.53 («0.001/7.17) | -377.35 («0.001/3.11) |
| **Mid-Temporal Gyrus, anterior div** | -92.30 («0.001/2.74) | -84.58 («0.001/1.04) |
| **Mid-Temporal Gyrus, posterior div** | -188.30 («0.001/3.08) | -171.46 («0.001/1.21) |
| **Middle Frontal Gyrus** | -250.07 («0.001/2.46) | -253.15 («0.001/1.05) |
| **Precentral Gyrus** | -310.07 («0.001/2.71) | -316.77 («0.001/1.37) |
| **Sup-Temporal Gyrus, anterior div** | -87.74 («0.001/3.01) | -84.65 («0.001/1.31) |
| **Sup-Temporal Gyrus, posterior div** | -129.47 («0.001/3.00) | -126.29 («0.001/1.40) |
| **Superior Frontal Gyrus** | -225.68 («0.001/1.80) | -224.18 («0.001/0.73) |
| **Temporal Pole** | -153.27 («0.001/1.82) | -145.58 («0.001/0.65) |
| **Angular Gyrus** | -192.6 («0.001/3.23) | -197.08 («0.001/1.69) |
| **Inf-Temporal Gyrus, anterior div** | -84.65 («0.001/2.51) | -73.97 («0.001/0.89) |
| **Inf-Temporal Gyrus, posterior div** | -121.68 («0.001/2.07) | -111.64 («0.001/0.77) |
| **Inf-Temporal Gyrus, temp-occipital part** | -128.48 («0.001/2.71) | -123.37 («0.001/1.09) |
| **Intracalcarine Cortex** | -459.57 («0.001/6.38) | -201.14 («0.001/2.09) |
| **Lat-Occipital Cortex, inferior div** | -190.45 («0.001/2.19) | -157.99 («0.001/0.69) |
| **Lat-Occipital Cortex, superior div** | -245.35 («0.001/2.02) | -231.68 («0.001/1.16) |
| **Mid-Temporal Gyrus, temp-occipital part** | -181.70 («0.001/3.32) | -175.84 («0.001/1.48) |
| **Postcentral Gyrus** | -228.78 («0.001/2.48) | -223.76 («0.001/1.39) |
| **Sup-Parietal Lobule** | -179.50 («0.001/2.64) | -178.27 («0.001/1.81) |
| **Supramarginal Gyrus, anterior div** | -143.52 («0.001/3.01) | -145.01 («0.001/1.50) |
| **Supramarginal Gyrus, posterior div** | -180.00 («0.001/2.94) | -190.31 («0.001/1.61) |
| **Cingulate Gyrus, anterior div** | -635.73 («0.001/8.52) | -476.77 («0.001/4.06) |
| **Cingulate Gyrus, posterior div** | -309.96 («0.001/4.56) | -320.57 («0.001/2.71) |
| **Cuneal Cortex** | -183.40 («0.001/4.07) | -152.85 («0.001/2.12) |
| **Frontal Medial Cortex** | -160.41 («0.001/3.92) | -104.09 («0.001/0.97) |
| **Frontal Orbital Cortex** | -194.45 («0.001/2.81) | -178.63 («0.001/0.91) |
| **Juxtapositional Lobule Cortex** | -169.78 («0.001/2.89) | -171.76 («0.001/1.34) |
| **Lingual Gyrus** | -319.14 («0.001/4.60) | -160.94 («0.001/1.41) |
| **Paracingulate Gyrus** | -394.22 («0.001/5.52) | -339.11 («0.001/1.69) |
| **Parahippocampal Gyrus, anterior div** | -139.13 («0.001/2.58) | -132.32 («0.001/0.94) |
| **Parahippocampal Gyrus, posterior div** | -124.81 («0.001/3.19) | -121.46 («0.001/1.30) |
| **Precuneous Cortex** | -339.96 («0.001/4.22) | -346.43 («0.001/2.71) |
| **Subcallosal Cortex** | -145.39 («0.001/3.22) | -128.58 («0.001/0.98) |
| **Central Opercular Cortex** | -454.89 («0.001/8.48) | -418.34 («0.001/4.36) |
| **Frontal Operculum Cortex** | -243.46 («0.001/7.62) | -242.79 («0.001/3.15) |
| **Heschls Gyrus** | -488.41 («0.001/12.06) | -293.71 («0.001/6.27) |
| **Occipital Fusiform Gyrus** | -380.06 («0.001/4.38) | -136.75 («0.001/0.96) |
| **Occipital Pole** | -132.12 («0.001/1.48) | -107.48 («0.001/0.59) |
| **Parietal Operculum Cortex** | -600.96 («0.001/12.22) | -476.64 («0.001/7.34) |
| **Planum Polare** | -211.32 («0.001/5.45) | -219.71 («0.001/2.44) |
| **Planum Temporale** | -388.60 («0.001/8.22) | -345.87 («0.001/4.67) |
| **Supracalcarine Cortex** | -303.88 («0.001/6.03) | -82.39 («0.001/2.06) |
| **Temp-Fusiform Cortex, anterior div** | -82.64 («0.001/2.41) | -76.43 («0.001/0.93) |
| **Temp-Fusiform Cortex, posterior div** | -145.43 («0.001/2.92) | -133.17 («0.001/1.08) |
| **Temp-Occipital Fusiform Cortex** | -444.47 («0.001/7.58) | -266.05 («0.001/2.91) |

Table S3: Welch's t-test statistic (p-value/Cohen's d) values for different cortical ROIs from the Oxford-Harvard atlas. For Cohen's d, higher values indicate a positive effect size for the first disease group specified. Values of above absolute value 0.2 are regarded as having a significant effect.

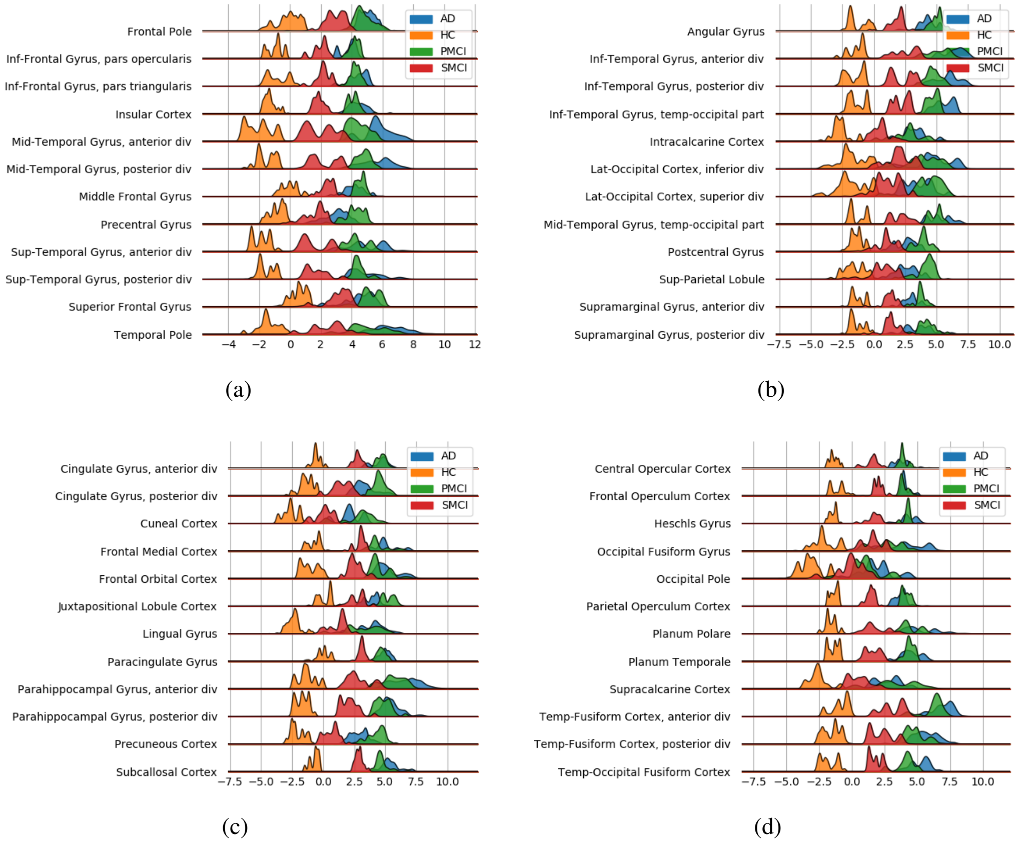


Figure S6: Cortical ROI-based difference in voxel level Brain-PAD scores averaged across subjects from clinical groups from OASIS3.

### G Stability of clinical results depending on dataset

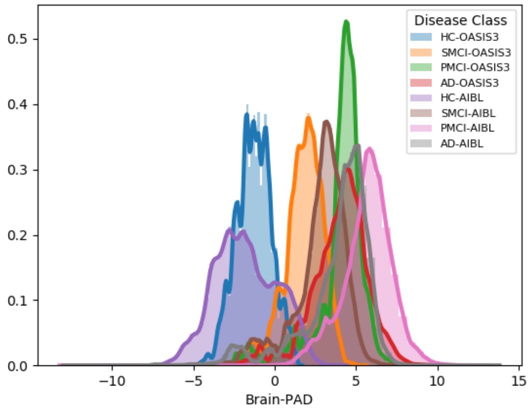


Figure S7: Histogram at voxel-level of Brain-PAD scores of certain clinical groups from OASIS3 and AIBL. Brain-PAD after applying the bias-adjustment scheme is calculated for every voxel and then aggregated to the mean across all subjects. Histograms in the plot are composed of the mean Brain-PAD values for all voxels in the brain.

In this section we explore whether LocalBrainAge is able to detect similar pathological changes in structural brain matter between subjects with varying degrees of cognitive impairment stemming from two different datasets, respectively OASIS3 and AIBL.

From Figure S7 we can infer that the effect size of the different groups with cognitive impairment are relatively similar between the two datasets, with the one noticeable difference that for AIBL, pMCI subjects tend to have higher local brain-PAD scores compared to AD subjects from the same study.

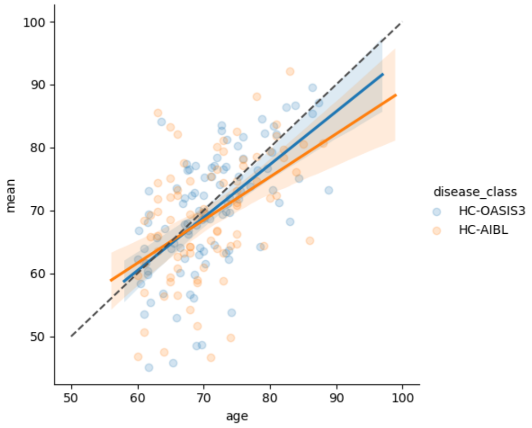


(a)

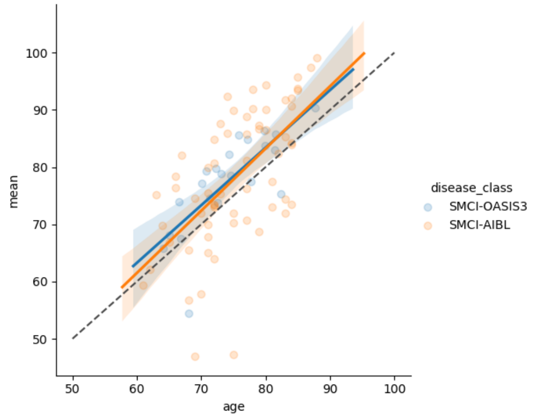


(b)

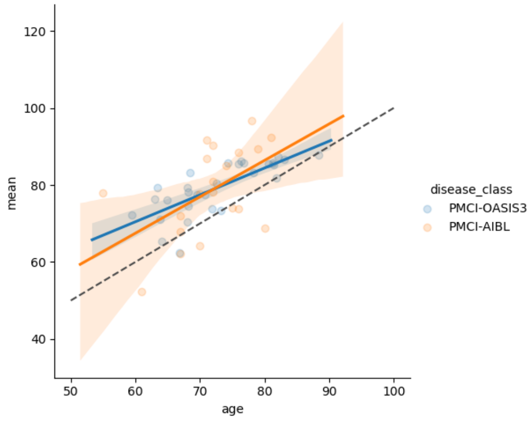


(c)

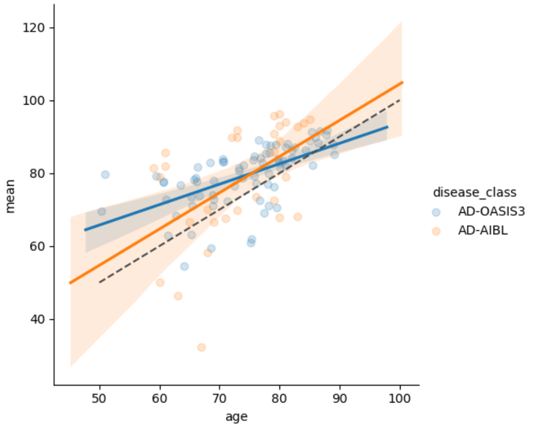


(d)

Figure S8: Scatter plots of chronological age versus averaged across all voxels predictive mean.

Looking from a whole-brain perspective, the patterns for the various disease groups are almost identical for OASIS3 and AIBL (Figure S8), irrespective of disease group.
